# Engineering the Mechanical Stability of a Therapeutic Affibody/PD-L1 Complex by Anchor Point Selection

**DOI:** 10.1101/2024.05.21.595133

**Authors:** Byeongseon Yang, Diego E. B. Gomes, Zhaowei Liu, Mariana Sá Santos, Jiajun Li, Rafael C. Bernardi, Michael A. Nash

## Abstract

Protein-protein complexes can vary in mechanical stability depending on the direction from which force is applied. Here we investigated the anisotropic mechanical stability of a molecular complex between a therapeutic non-immunoglobulin scaffold called Affibody and the extracellular domain of the immune checkpoint protein PD-L1. We used a combination of single-molecule AFM force spectroscopy (AFM-SMFS) with bioorthogonal clickable peptide handles, shear stress bead adhesion assays, molecular modeling, and steered molecular dynamics (SMD) simulations to understand the pulling point dependency of mechanostability of the Affibody:(PD-L1) complex. We observed diverse mechanical responses depending on the anchor point. For example, pulling from residue #22 on Affibody generated an intermediate unfolding event attributed to partial unfolding of PD-L1, while pulling from Affibody’s N-terminus generated force-activated catch bond behavior. We found that pulling from residue #22 or #47 on Affibody generated the highest rupture forces, with the complex breaking at up to ∼ 190 pN under loading rates of ∼10^4^-10^5^ pN/sec, representing a ∼4-fold increase in mechanostability as compared with low force N-terminal pulling. SMD simulations provided consistent tendencies in rupture forces, and through visualization of force propagation networks provided mechanistic insights. These results demonstrate how mechanostability of therapeutic protein-protein interfaces can be controlled by informed selection of anchor points within molecules, with implications for optimal bioconjugation strategies in drug delivery vehicles.

## Introduction

Protein-protein complexes are inherently anisotropic objects, with non-covalent interactions that stabilize the structure oriented variably in different directions. When tension is applied to such a system, the amount of force that the complex can resist before dissociating will significantly depend on the anchor points (i.e. amino acid residues) through which tension is applied. This concept, known as mechanical anisotropy, has been demonstrated on folded protein domains^1–5^, nucleic acids^6^ and protein-protein complexes^7–11^, among others^12^. Although this conceptual framework has existed in the molecular biophysics community for years, its potential for enhancing binding strength for applications in drug delivery is less explored. In particular, for nanoparticle and cell-based therapies, binding proteins attached to the particle or cell surface to achieve specific targeting are subjected to hydrodynamic shear forces inside the body. If these binding proteins have been optimized for high affinity interactions at equilibrium (i.e. in the absence of force), they can perform poorly and unbind at low forces when exposed to shear stress.^13,14^ Therefore, in developing drug delivery systems for operation under conditions of hydrodynamic flow, the concept of imparting mechanical stability through anchor point selection to improve interaction and adhesion strength is of significant interest.

Non-immunoglobulin (non-Ig) scaffolds have significant potential as next-generation therapeutics due to their small size and ease of production.^15,16^ The Affibody (AFF) scaffold based on the Z-domain of *S. aureus* protein A in particular is a promising alternative scaffold comprising three bundled α-helices.^17–19^ Solvent accessible residues contained within α-helices 1 and 2 of the AFF scaffold have been mutagenized to isolate AFF variants with specific binding activity towards several therapeutic drug targets, for example, programmed cell death ligand 1 (PD-L1),^20–22^ an immune checkpoint protein and major target in cancer immunotherapy.^23–26^ In fact, an AFF variant that binds PD-L1 with high affinity was reported as a potential candidate for *in vivo* tumor imaging with good specificity and rapid clearance.^20,21^

AFM-based single-molecule force spectroscopy (AFM-SMFS) is a powerful technique to study the mechanical responses of protein interactions under force.^27,28^ Single protein-protein complexes can be dissociated under mechanical tension to study a variety of mechanical responses, such as classical slip bonds, force-activated catch bonds, intermediate unfolding states, and other structural transitions.^29–33^ Recently we reported on a protein bioconjugation and surface chemical approach relying on non-canonical amino acid (NCAA) incorporation and click chemistry to install peptide handles onto single side chains, and apply tension at any position within a protein sequence.^7^ The peptide handle^10,11,34^ called Fgβ served as an orthogonal pulling point within the sequence that could be recognized by an SdrG domain attached to a cantilever tip, allowing us to study the anchor point dependence of mechanostability in protein-protein complexes.

Here, using AFM-SMFS we analyzed the mechanical stability of the AFF:(PD-L1) complex when pulled from five different anchor points on AFF. Combined with in silico SMFS, which is based on steered molecular dynamics (SMD) simulations, we observed different mechanical responses and changes in the unbinding pathway depending on the pulling point. We further compared the most and least stable pulling points using a bead-based shear stress adhesion assay^35^ to analyze how these single-molecule mechanical properties manifest under conditions of multivalent collective binding under the influence of shear forces.^35– 37^ We found that in particular, pulling AFF from central positions within the sequence generated higher resistance to mechanical load (i.e. higher rupture forces), which resulted in enhanced bead adhesion under shear flow. The SMD simulations provide crucial information about the mechanical properties of the (PD-L1):AFF complex, confirming the mechanical anisotropy and revealing differences in unfolding behavior, rupture forces, for propagation pathways and contact interactions depending on the pulling points.

## Results

### Selection of Pulling Points, Protein Preparation, and AFM Measurement Setup

An AFF variant that binds PD-L1 was previously engineered by random mutagenesis and screening of solvent-accessible residue variants within the first two α-helices of AFF.^20,21^ In our study, recombinant AFF and the extracellular domain of human PD-L1 (PD-L1-ECD) were prepared by overexpression in *E. coli* to first confirm binding (**Figure S1** and **Figure S2**).^38^ Since AFF is only 59 amino acids long and consists of three α-helices, there were limited positions within the sequence where we could install orthogonal pulling points while minimizing perturbations to the binding epitope. Analyzing structural models of AFF, we determined that potential positions for pulling point insertion would be at: 1) the N-terminus; 2) within the loop between α1 and α2 helices; 3) within the loop between α2 and α3 helices; 4) solvent accessible residues on α3 helix; and 5) at the C-terminus (**Figure 1A, 1B**). Based on these considerations, five pulling points within AFF were chosen by selecting one residue from each of these regions. The chosen pulling points on AFF were the N-terminus (M1), C-terminus (G60) and three internal sequence positions (N22, S40, S47; **Figure 1B**). In all experiments, the PD-L1 extracellular domain was always anchored through its C-terminus to mimic its orientation on the cell surface.

**Figure 1.**
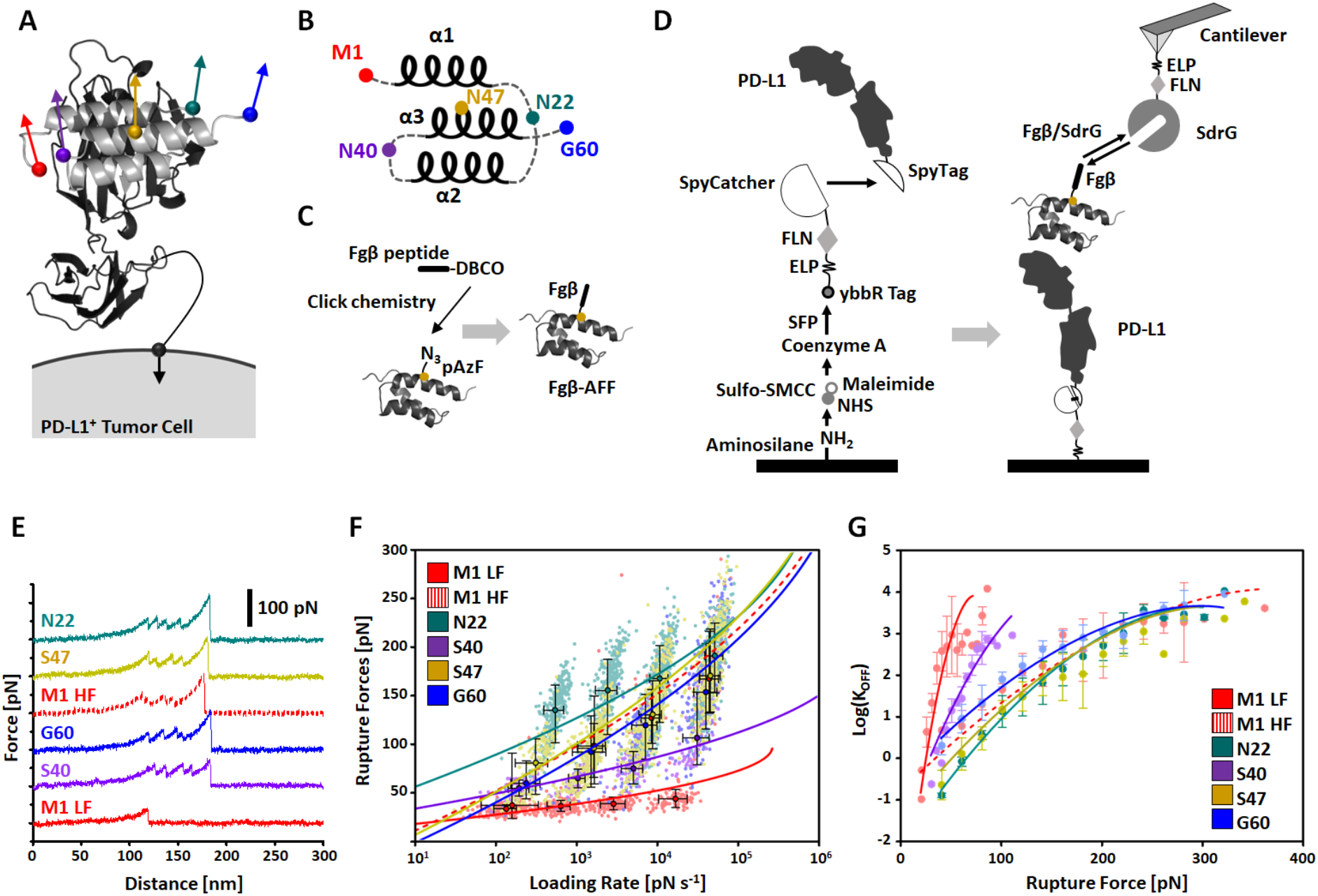
Pulling point selection and mechanostability of the (PD-L1):AFF complex analyzed by single-molecule AFM. (A) Prediction of AFF structure in complex with PD-L1 by AlphaFold. Pulling points on AFF are indicated as colored spheres. The extracellular domain of PD-L1 is fixed at the C-terminus according to the natural anchor geometry on the cell surface. (B) AFF consists of three α helices connected by short flexible loops. The pulling points were selected from the termini, loops connecting α helices, and in the middle of the α3 helix. (C) Bioorthogonal conjugation of a fibrinogen β (Fgβ) peptide to AFF via non-canonical amino acid incorporation and click chemistry. P-azido-phenylalanine was introduced at a single desired pulling point using amber suppression and clicked to a synthetic Fgβ-DBCO peptide. (D) Schematic illustration of the surface chemistry, site-specific protein immobilization, and AFM setup with freely diffusing receptor system. (E) Typical AFM force−extension traces of the (PD-L1):AFF complex rupture traces under five different pulling geometries, including M1 high force population (M1 HF; Dashed line) and M1 low force population (M1 LF; Solid line). Unfolding of two FLN fingerprint domains was used to filter the curves for specific single-molecule interactions. For the N-terminal (low force) pulling points, the (PD-L1):AFF interaction was not sufficiently strong to unfold the FLN fingerprint domains. (F) Dynamic force spectra of (PD-L1):AFF complex rupture events. Black-lined circles represent the median rupture force/loading rate at each pulling speed of 0.1×10^3^, 0.4×10^3^, 1.6×10^3^, and 6.4×10^3^ nm s^−1^. Error bars are ±1 s.d. Solid and dashed lines are least square fits to the Dudko-Hummer-Szabo (DHS) model. (G) The force-dependent off rate of the (PD-L1):AFF complex was plotted against rupture force and fitted to DHS model extract Δx, *k*_*0*_ and ΔG.

To dissociate the (PD-L1):AFF complex by applying tension through a given pulling point, we used NCAA incorporation into AFF combined with click chemistry to install a peptide pulling handle at a given position.^28^ We first introduced an Amber stop codon into the gene cassette encoding AFF at the desired position. We then employed site-specific incorporation of p-Azido-L-phenylalanine (pAzF) by amber suppression in *E. coli*, and confirmed incorporation by high resolution mass spectrometry (HR-MS) (**Figure S3**). To attach the Fgβ peptide as a pulling handle, a synthetic Fgβ peptide containing a C-terminal dibenzocyclooctyne (DBCO) group was conjugated onto the desired position of AFF using a copper-free click reaction between the azide group of pAzF and the DBCO group on Fgβ (**Figure 1C**). Following conjugation, we characterized Fgβ-AFF conjugates using SDS-PAGE and HR-MS analyses (**Figure S3**). For the N-terminal pulling points, AFF was separately prepared as a fusion with an N-terminal Fgβ peptide sequence (**Figure S4**). Each AFF pulling point variant was separately cloned with a single amber codon at the given position, produced in *E. coli*, conjugated with Fgβ and measured in separate AFM-SMFS experiments.

For the AFM experimental setup, we produced the PD-L1 extracellular domain with a C-terminal SpyTag (PD-L1-SpyTag) (**Figure S5**). As PD-L1 is naturally a transmembrane protein with C-terminal transmembrane domain, SpyTag was fused to the C-terminal side of the recombinant extracellular PD-L1 domain to mimic the natural anchor point of PD-L1 in the membrane. The SpyTag was then further conjugated to a SpyCatcher domains *via* spontaneous isopeptide bond formation.^39,40^ SpyCatcher was produced as polyprotein (SpyCatcher-FLN-ELP-ybbR) where the fourth domain of Dictyostelium discoideum F-actin cross-linking filamin (FLN) served as an SMFS fingerprint, an elastin-like polypeptide (ELP) sequence served as a flexible linker, and a ybbR tag facilitated site-specific covalent surface immobilization (**Figure S5**).^39–41^ PD-L1-SpyTag was conjugated to SpyCatcher-FLN-ELP-ybbR and further site-specifically and covalently immobilized onto coenzyme A (CoA)-functionalized coverglass substrates *via* ligation by 4′-phosphopantetheinyl transferase (SFP) (**Figure 1D** and **Figure S5**). For the free diffusion measurement setup, SD-repeat protein G (SdrG) from *S. epidermidis* was prepared as a fusion protein of the form SdrG-FLN-ELP-ybbR, and immobilized onto silicon nitride cantilever tips *via the* ybbR tag. Cantilevers functionalized in this manner could be used to bind Fgβ-AFFs (**Figure 1D**) and dissociate them from surface-immobilized PD-L1 by applying tension through the various pulling points.^7^ SdrG binds Fgβ with moderate equilibrium binding affinity (K_D_ ∼400 nM) which enabled rapid exchange of Fgβ-conjugated molecules on the cantilever tip. This measurement approach is able to constantly probe fresh molecules on both the surface (by actuating the AFM X-Y stage) and cantilever (by reversible ligand dissociation/exchange), thereby avoiding irreversible protein unfolding.^7,28,34,42^ Since the SdrG:Fgβ complex can withstand up to ∼2 nN of force prior to rupture, it is capable of breaking the (PD-L1):AFF binding interface. With this AFM setup, (PD-L1):AFF complexes with five different pulling geometry were probed at four different pulling speeds (0.1×10^3^, 0.4×10^3^, 1.6×10^3^, 6.4×10^3^ nm s^−1^) using constant speed AFM-SMFS. All data traces were filtered for two 34 nm contour length increments which originated from the two FLN fingerprint domains, one from the SdrG bound to the cantilever, and one from the PD-L1 bound to the surface. This fingerprint filtering method allowed us to identify single-molecule level interactions (**Figure 1E**) in large datasets consisting of tens of thousands of AFM force-extension curves. FLN unfolding was followed by rupture of (PD-L1):AFF complex. In some cases, we found that the (PD-L1):AFF complex rupture force range was lower than the unfolding force of FLN. Therefore, data traces with no FLN unfolding were also analyzed in the case of the N-terminal pulling point (M1, red) (**Figure 1E**).

### Alteration of Pulling Geometry Drastically Changes the Unbinding Energy Landscape

We analyzed the (PD-L1):AFF complex rupture events under different pulling geometries. In total we conducted ten different AFM measurement trials using two replicates of five pulling points, collecting 109,067 total AFM-SMFS force-extension traces in total. Of these, 33,365 curves were selected for further analysis, and finally the contour length and the quality filters were applied generating 4,868 rupture force data points. Final data points were plotted as a function of the loading rate (**Figure 1F**), demonstrating how the mechanostability of the (PD-L1):AFF complex strongly depends on the pulling point. Depending on the anchor point residue on AFF, different characteristics were observed in terms of rupture force ranges, loading rate dependencies, the presence of multiple unbinding pathways, and energy profile parameters (**Figure 1F** and **1G**), as discussed in detail below.

For pulling point M1, the rupture force range was the lowest at all pulling speeds compared to the other pulling geometries, ranging from 34 ± 4 pN at 0.1 × 10^3^ nm s^−1^ to 44 ± 9 pN at 6.4 × 10^3^ nm s^−1^ (**Table 1**). Interestingly, we observed a new rupture force population as pulling speed increased. This new population showed much higher rupture forces, more similar to the other pulling points such as S47 and G60 and with similar loading rate dependency. The ratio of this new high force population (M1 HF) to low force population (M1 LF) increased as the pulling speed increased, showing catch bond-like behavior (**Figure S6**) as manifested in a constant speed pulling protocol.^31,43^

**Table 1.** Energy landscape parameters of the (PD-L1):AFF complex rupture under different pulling geometries by AFM-SMFS with Bell-Evans (BE) and Dudko-Hummer-Szabo (DHS) model fitting and the dissociation constant values measured by beads-based equilibrium binding affinity analysis via flow cytometry. ^a^HF: High force population, ^b^PU: Partial unfolding.

| Pulling Points | $\Delta x$ [nm] (BE) | $\Delta x$ [nm] (DHS) | $\ln(k_{off})$ (BE) | $\ln(k_{off})$ (DHS) | $\Delta G$ [$k_B T$] (DHS) | Rupture force [pN] $0.1 \times 10^3$ nm s<sup>-1</sup> | Rupture force [pN] $6.4 \times 10^3$ nm s<sup>-1</sup> | $K_D$ [nM] |
| --- | --- | --- | --- | --- | --- | --- | --- | --- |
| **M1** | $2.0 \pm 0.2$ | $1.3 \pm 0.2$ | $-11.8 \pm 1.8$ | $-4.5 \pm 1.8$ | $15.3 \pm 2.0$ | $34 \pm 4$ | $44 \pm 9$ | $28.1 \pm 13.8$ |
| HF<sup>a</sup> | $0.18 \pm 0.00$ | $0.25 \pm 0.03$ | $-0.2 \pm 0.2$ | $-1.7 \pm 0.8$ | $13.0 \pm 0.9$ | $37 \pm 34$ | $167 \pm 43$ | |
| **N22** | $0.35 \pm 0.03$ | $0.36 \pm 0.03$ | $-7.6 \pm 1.0$ | $-5.2 \pm 0.7$ | $15.8 \pm 0.8$ | $135 \pm 30$ | $191 \pm 36$ | $22.3 \pm 4.5$ |
| PU<sup>b</sup> | $0.87 \pm 0.05$ | $1.0 \pm 0.1$ | $-16.6 \pm 1.3$ | $-13.3 \pm 1.5$ | $25.1 \pm 1.5$ | $100 \pm 14$ | $121 \pm 16$ | |
| **S40** | $0.65 \pm 0.02$ | $0.78 \pm 0.25$ | $-5.2 \pm 0.3$ | $-5.7 \pm 1.8$ | $18.2 \pm 4.0$ | $55 \pm 8$ | $106 \pm 37$ | $22.5 \pm 7.8$ |
| **S47** | $0.23 \pm 0.03$ | $0.23 \pm 0.02$ | $-1.2 \pm 0.8$ | $-1.5 \pm 0.6$ | $12.6 \pm 0.5$ | $81 \pm 28$ | $170 \pm 42$ | $35.5 \pm 7.5$ |
| **G60** | $0.22 \pm 0.01$ | $0.24 \pm 0.02$ | $-0.6 \pm 0.1$ | $-1.0 \pm 0.5$ | $12.3 \pm 0.4$ | $60 \pm 19$ | $154 \pm 44$ | $37.1 \pm 11.3$ |

Meanwhile, the pulling point N22 generated the most mechanostable (PD-L1):AFF interaction, with the highest rupture forces across all pulling speeds from 135 ± 30 pN at 0.1 × 10^3^ nm s^−1^ to 191 ± 36 pN at 6.4 × 10^3^ nm s^−1^ (**Table 1**). This force range was ∼4.0-4.3 fold higher than the low force population found for pulling point M1. Pulling point N22 was furthermore characterized by an alternative unbinding pathway with an intermediate unfolding state that was not found in any other pulling geometry. We regularly detected an additional intermediate unfolding peak prior to final rupture for pulling point N22 (**Figure S7**). The contour length increment associated with this unfolding event was found to be ∼14 nm. Assuming ∼0.35 nm / amino acid in contour length space, this corresponds to an unfolding event of ∼40 amino acids. If we assigned this unfolding event to a structural transition within AFF, it would correspond to unfolding of ∼68% of the AFF length while maintaining strong binding to PD-L1 (**Figure S7**). We judged this scenario unlikely, and therefore assigned the intermediate unfolding increment observed for the N22 pulling point to partial unfolding of PD-L1. Therefore, we concluded that mechanostability of the (PD-L1):AFF complex when pulled from point N22 was significantly high such that force applied to AFF can partially unfold PD-L1 prior to unbinding from PD-L1. It is important to note here that our recombinant PD-L1 consists of two separate domains that could conceivably (un)fold independently from one another.

For pulling points S40, S47, and G60, the (PD-L1):AFF complexes showed typical unimodal behavior without intermediate unfolding events or catch bond behavior. While S40 exhibited low to moderate rupture forces, ranging from 55 ± 8 pN at 0.1 × 10^3^ nm s^−1^ to 106 ± 37 pN at 6.4 × 10^3^ nm s^−1^ (**Table 1**), both S47 and G60 showed low to high force rupture rupture events with a steep loading rate dependency. At a slow pulling speed of 0.1 × 10^3^ nm s^−1^, weak interactions were observed with a rupture force of 81 ± 28 pN and 60 ± 19 pN for S47 and G60, respectively, however at the high pulling speed of 6.4 × 10^3^ nm s^−1^, strong interactions were observed, as much as 170 ± 42 pN and 154 ± 44 pN for S47 and G60, respectively.

Unbinding energy landscape parameters generated by the various pulling geometries were estimated by parameter fitting using Bell-Evans (BE)^44,45^ and Dudko-Hummer-Szabo (DHS)^46^ models. The relevant parameters were the distance to the energy barrier (Δx) and the off-rate (*k*_*off*_) for the BE model (**Figure 1F**). For the DHS model, we obtained the energy barrier heights (ΔG), Δx and *k*_*off*_ in order to generate a force-dependent off-rate plot (**Figure 1G**). The estimated parameters from both models are listed in **Table**

**1**. We note that these models are highly simplified and assume a simple two-state system, which is not the case for our complex as the intermediate events for pulling point N22 and catch bond behavior for pulling point M1 demonstrate. Furthermore, differences in the fitted zero-force *k*_*off*_ are spurious since for our system all anchor point variants have the same equilibrium binding affinity (see below) and presumably very similar equilibrium off-rates. Nonetheless, the model fitting demonstrated that unbinding of the (PD-L1):AFF complex from different pulling points exhibits different energy profiles, with fitted Δx parameters varying up to ∼9-fold and ΔG varying up to ∼1.9-fold depending on the pulling point.

### Minimal Dependency of Conjugation Points on Equilibrium Binding Affinity

To investigate the effect of anchor point engineering on equilibrium binding behavior, we performed beads-based flow cytometric binding assays to measure the equilibrium dissociation constant (K_D_) between (PD-L1):AFF using a fluorescent dye conjugated to the bioorthogonal side chain used for pulling in the AFM experiments. AFF variants were prepared where FAM-DBCO was conjugated to each of five pulling points (instead of Fgβ) using the same click reaction. PD-L1 was immobilized on amine-functionalized polystyrene (PS) beads in the same way for AFM setup (**Figure 2A**). The AFF-FAM conjugates were titrated across a concentration range and the fraction of fluorescently labeled beads was analyzed by flow cytometry to extract fitted K_D_ values (**Figure 2B**) for each AFF variant. We found K_D_ values to be 28.1 ± 13.8 nM, 22.3 ± 4.5 nM, 22.5 ± 7.8 nM, 35.5 ± 7.5 nM, and 37.1 ± 11.3 nM for AFF variants M1, N22, S40, S47, and G60, respectively (**Figure 2B**). Despite the quantitative and qualitative differences in unbinding behavior observed for the various pulling points in the AFM experiments, the equilibrium analysis revealed that binding affinity was not significantly changed by introduction of the bioorthogonal azide group at the respective position. This also validated the rationale for pulling point selection, where we targeted only flexible loops, flexible termini or solvent exposed side chains in AFF located away from the binding interface with PD-L1.

**Figure 2.**
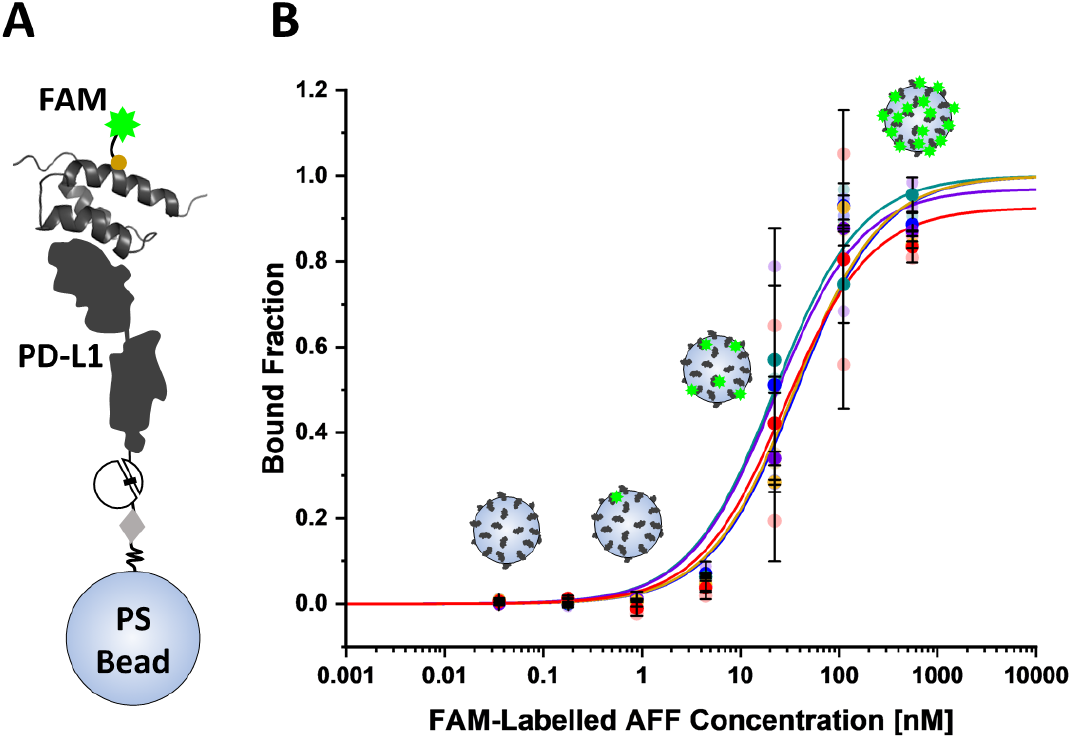
Bead-based equilibrium binding affinity analysis via flow cytometry. (A) Schematic illustration of surface chemistry, site-specific protein immobilization, and fluorescent dye conjugation. (B) Binding curves and the fitting to determine binding affinity of the (PD-L1):AFFibody complex with different conjugation points. Error bars are ±1 s.d.

### Mechanical Insight of (PD-L1):AFF Complex Rupture by SMD

Next we used Steered Molecular Dynamics (SMD) to simulate the (PD-L1):AFF system while applying tension through the same pulling points as in the AFM-SMFS experiments (**Figure 1A**). Each geometry was probed at three constant pulling speeds by SMD (2.5 × 10^7^, 2.5 × 10^8^, and 2.5 × 10^9^ nm s^−1^). We examined and compared the unfolding and rupture events, and plotted against the loading rate together with the experimental dataset on a single set of axes (**Figure 3A**).^47^ Consistent with the AFM-SMFS measurements, our SMD simulations demonstrated significant mechanical anisotropy, with different pulling geometries exhibiting distinct characteristics in terms of protein unfolding, rupture force ranges, loading rate dependencies, unbinding pathways, and force profiles.

**Figure 3.**
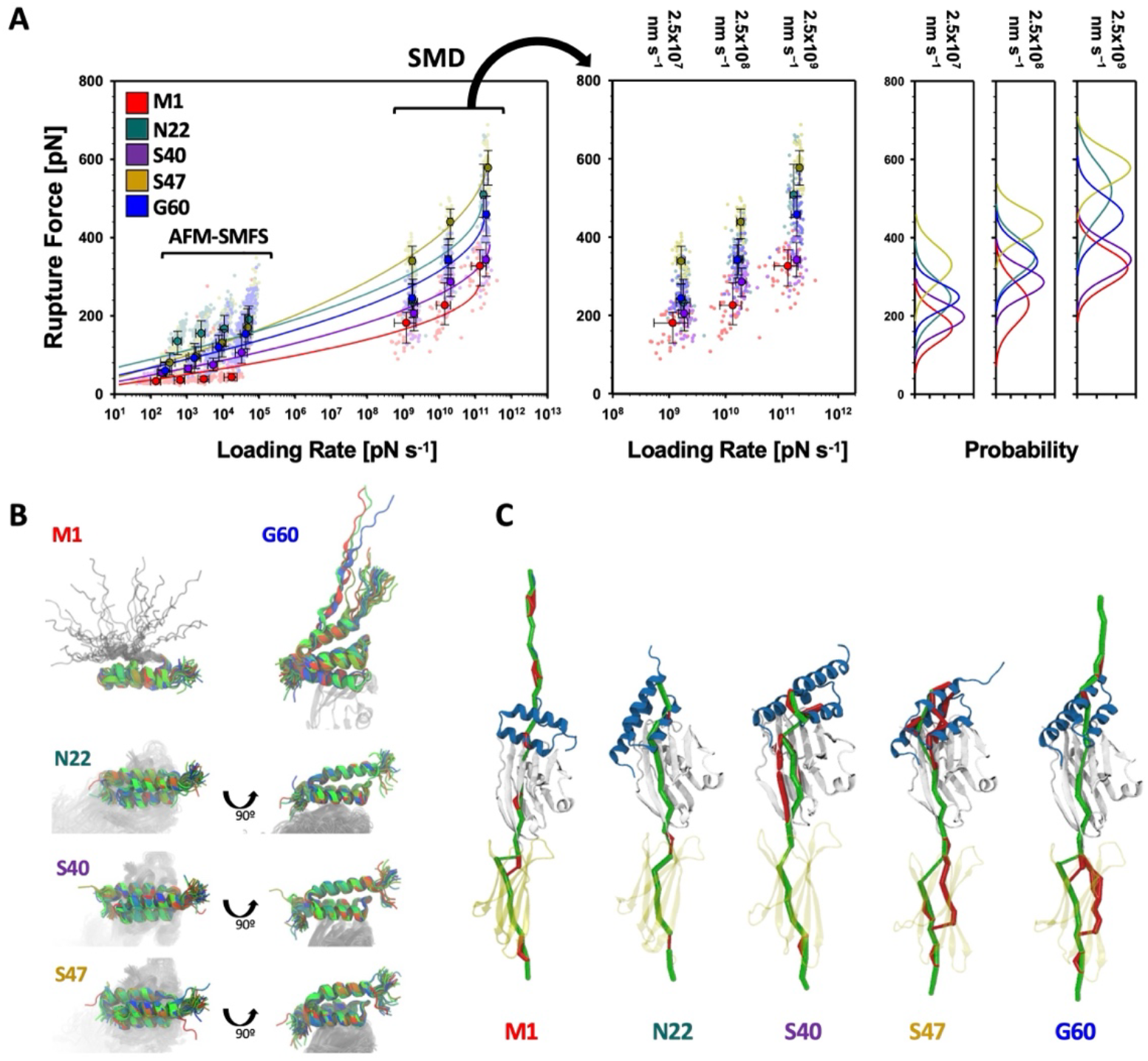
Steered molecular dynamics simulations of pulling (PD-L1):AFFibody complex from different anchor points. (A) Combined experimental and simulated dynamic force spectra and the histogram of the (PD-L1):AFFibody complex rupture. Black-lined circles represent the median rupture force/loading rate at each pulling speed. Error bars are ±1 s.d. Solid lines in dynamic force spectra are least square fits to the DHS model. Solid lines in histogram are Gaussian fit. (B) The closest trajectory frame to the peak force measured from all replicas at 2.5×10^7^ nm s^−1^ after fitting to AFF’s backbone atoms from residues 5 to 55. When pulling from the N-terminal, fitting was limited to residues 25 to 55 due to unfolding of α1. C and N terminals have 40 replicas each, while N22, S40 and S47 have 48 replicas each. (C) Predominant force propagation pathways. For most replicas and pulling points (except C-terminal) the optimal force propagation path (green) from the pulling residue to the anchor exits the AFF through Asp36_AFF_, passing through Arg96_PD-L1_, and then propagating up to Gly93_AFF_ and crossing to PD-L1 through Lys112_PD-L1_ or Val113_PD-L1_. Suboptimal paths (red) less frequently take alternative routes through the interface. While pulling from Ser47_AFF_, most replicas commonly showed two suboptimal pathways that may reinforce the complex and explain its higher resilience. The width of the force propagation pathways is weighted by the correlation strength in the dynamic network.

Analysis of the (PD-L1):AFF simulated force traces revealed multiple peaks along the extension. A visual inspection of the SMD trajectories and of the nearest snapshot prior to rupture (**Figure 3B**) revealed sequential unfolding of the α1 helix when pulling from the N-terminus (**Video S1**). N-terminal pulling also resulted generally in the lowest rupture forces across all simulated pulling speeds (**Table 2**), and the broadest spread of loading rates for a given pulling speed (**Figure 3A**). When pulling from the C-terminus, we also regularly observed peeling of the α2 helix. Inspection of the forces and (PD-L1):AFF contact area for each pulling point reveals that for internal pulling residues (N22, S40, and S47), the peak force frequently coincided with complex rupture and loss of contact between the proteins. In the case of the N- and C-terminal pulling points (M1 and G60), single step breakage was less frequently observed, with the proteins exhibiting intermediate unfolding and remaining bound following the peak force (**Figure S9**).

**Table 2.** Energy landscape parameters of the (PD-L1):AFFibody complex rupture with different pulling geometries from SMD simulation with Bell-Evans (BE) model and Dudko-Hummer-Szabo (DHS) model fitting.

| Pulling Points | $\Delta x$ [nm]<br>(BE) | $\Delta x$ [nm]<br>(DHS) | $\ln(k_{\text{off}})$<br>(BE) | $\ln(k_{\text{off}})$<br>(DHS) | $\Delta G$ [k<sub>B</sub>T]<br>(DHS) | Rupture force [pN]<br>$2.5 \times 10^9 \text{ nm s}^{-1}$ |
| --- | --- | --- | --- | --- | --- | --- |
| **M1** | $0.31 \pm 0.04$ | $0.79 \pm 0.02$ | $1.8 \pm 1.4$ | $-5.7 \pm 1.8$ | $27.3 \pm 6.0$ | $328 \pm 46$ |
| **N22** | $0.28 \pm 0.07$ | $0.51 \pm 0.07$ | $-4.8 \pm 4.1$ | $-8.0 \pm 1.1$ | $31.2 \pm 6.4$ | $510 \pm 68$ |
| **S40** | $0.31 \pm 0.03$ | $0.60 \pm 0.07$ | $-0.4 \pm 1.3$ | $-4.4 \pm 1.6$ | $28.2 \pm 1.4$ | $344 \pm 44$ |
| **S47** | $0.18 \pm 0.02$ | $0.36 \pm 0.02$ | $-0.0 \pm 1.2$ | $-4.1 \pm 0.9$ | $27.3 \pm 0.8$ | $579 \pm 44$ |
| **G60** | $0.25 \pm 0.03$ | $0.53 \pm 0.06$ | $-0.8 \pm 1.8$ | $-6.1 \pm 2.0$ | $29.5 \pm 2.7$ | $459 \pm 51$ |

For pulling points N22, S40 and S47, the (PD-L1):AFF complex showed typical unimodal behavior without additional unfolding events or catch bond-like behavior. Pulling from N22 showed a stable interaction of the complex and generated the second highest rupture forces at all the simulated pulling speeds (e.g., 510 ± 68 nN at 2.5 × 10^7^ nm s^−1^ (**Table 2**)). These simulated rupture values were ∼1.5 fold higher than the low force population of the N_terminal_ pulling point. Pulling point N22 was not found in the simulations to generate partial unfolding of PD-L1 as was observed in the experiments (**Figure S7**). Similarly to AFM-SMFS, pulling from S40 showed a low to moderate range of ruptures (344 ± 44 nN at 2.5 × 10^7^ nm s^−1^), while S47 showed the highest rupture force range with strong loading rate dependency (579 ± 44 nN at 2.5 × 10^7^ nm s^−1^).

### Force propagation pathway analysis

We used generalized-correlation-based dynamical network analysis^48^ to extract correlations of motion from multiple (PD-L1):AFF simulation replicas for each pulling point and characterize the force propagation pathways^9^. For most replicas and pulling points (except C-terminal), the optimal force propagation pathways (**Figure 3C**, blue) between the AFF pulling point and the anchor point at PD-L1’s C-terminus exited AFF through residue Asp36_AFF_, passing through Arg96_PD-L1_, and propagating up to Gly93_AFF_ before crossing to PD-L1 through Lys112_PD-L1_ or Val113_PD-L1_. The suboptimal paths (**Figure 3C**, red) less frequently took alternative pathways across the binding interface. While pulling from Ser47_AFF_, most replicas showed two suboptimal pathways that may reinforce the complex and explain its higher force resilience. Aligned with the AFM-SMFS experiments, these findings explain the different rupture force populations observed as pulling speed increased. The simulations did not capture two distinct rupture force populations as was observed in AFM-SMFS (**Figure S6**).

### Enhanced microbead adhesion under shear stress

We next investigated how these anisotropic stability effects would influence microbead adhesion strength that arises from collective and multivalent interactions. We chose three internal pulling points for microbead adhesion assays based on the AFM-SMFS and SMD simulations: N22 (highest mechanostability by AFM-SMFS); S47 (highest mechanostability by SMD); and S40 (low mechanostability). Although M1 showed the lowest rupture force population in both AFM-SMFS and SMD simulations, it was not selected for the bead assays due to catch bond behavior found in the AFM-SMFS experiments (**Figure S6**).

We used the hydrodynamic shear-based spinning disk assay (SDA)^35,37^ where (PD-L1)-modified beads were deposited onto AFF-modified coverglass surfaces and exposed to a rotational hydrodynamic shear field. We designed and prepared a new Cys-SpyCatcher protein by introducing one cysteine in the middle of the short linker connecting FLN and SpyCatcher (**Figure S8**). Cys-SpyCatcher was DBCO-functionalized using DBCO-PEG_4_-maleimide and conjugated to AFF at the desired position via click chemistry (**Figure 4A** and **S8**). AFF-pAzF-DBCO-Cys-SpyCatcher for each of these three internal conjugation points was then immobilized onto amino-functionalized coverglass disks in the same way as was used in the AFM-SMFS setup. PD-L1 was immobilized onto PS beads in the same way as previously mentioned (**Figure 4A**). (PD-L1)-modified PS beads were adhered onto the AFF-immobilized coverglass (**Figure 4B**). As the disk was spun, beads experienced a gradient of shear stress that increased from the disk center to the edge. The higher shear stress at the outer edge of the disk disrupted the (PD-L1):AFF interactions, resulting in loss of bead adhesion. The sigmoidal decrease in bead density as a function of the distance from the disk center was plotted and analyzed to determine τ50, a parameter that represents the amount of shear stress required to detach 50% of the beads.

**Figure 4.**
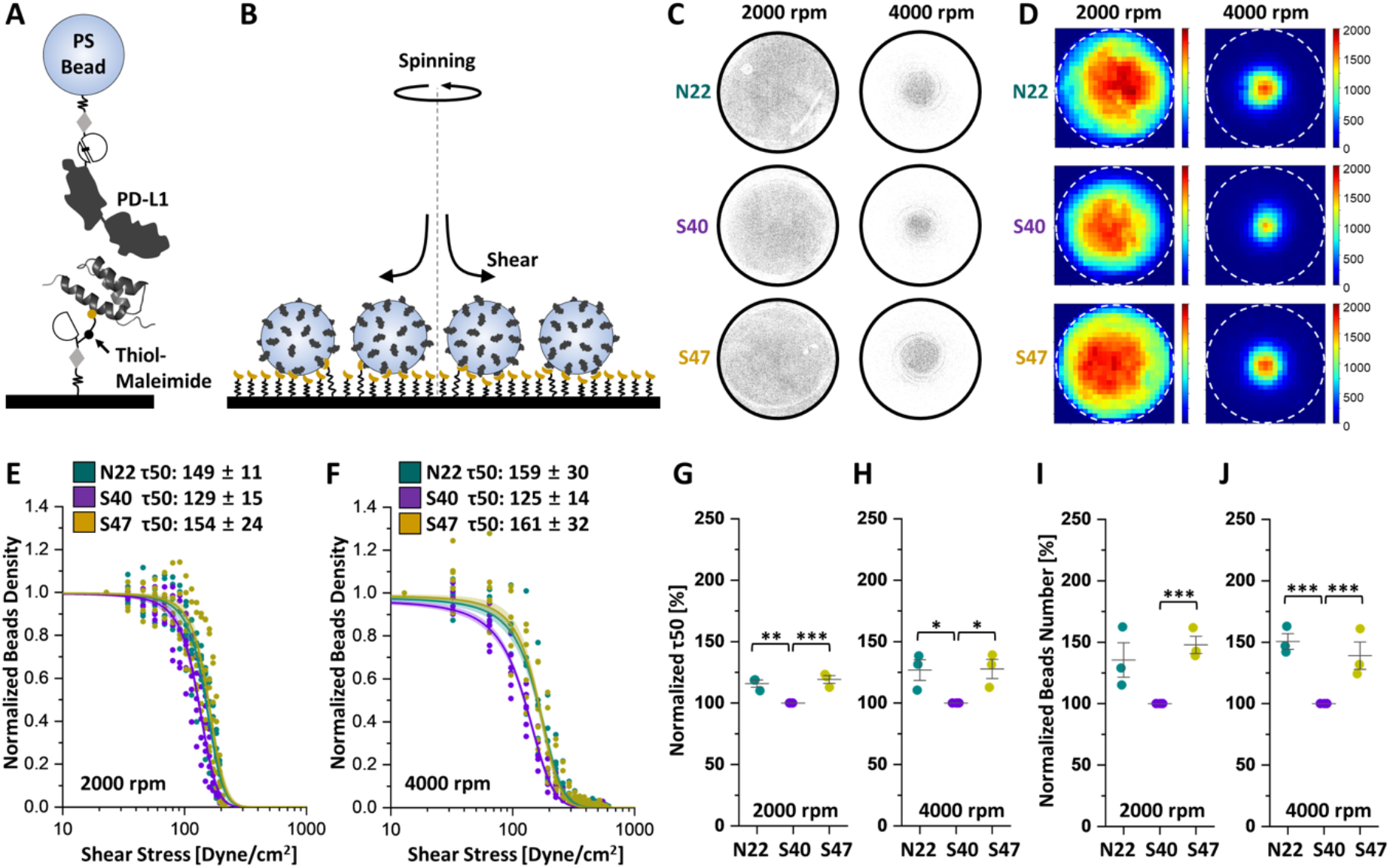
Analysis of microbead adhesion under shear flow mediated by AFF:(PD-L1) complexes. (A) Schematic illustration of surface chemistry, site-specific protein conjugation, and immobilization. (B) PS beads coated with PD-L1 are adhered to an AFF-modified glass disk and exposed to a shear stress generated by spinning. (C) Typical raw images and (D) beads density map of AFF-modified disks with three different AFF anchor points following spinning at 2000 or 4000 rpm. Normalized bead density plots *vs*. shear stress for the three different AFF anchor points at spinning speeds of (E) 2000 and (F) 4000 rpm . Plotted data from six glass disks were fitted with a sigmoid model to extract τ50, the shear stress required to detach half the bead population. Comparison of normalized τ50 of the AFF:(PD-L1) complex with three different conjugation points at the spinning speeds of (G) 2000 and (H) 4000 rpm. Comparison of normalized total beads number of the AFF:(PD-L1) complex with three different conjugation points at the spinning speeds of (I) 2000 and (J) 4000 rpm. Shades indicate 95% confidence intervals from fitting. Teal: AFF-N22pAzF, Purple: AFF-S40pAzF, and Dark yellow: AFF-S47pAzF. n.s. p ≥ 0.05; *p < 0.05; **p < 0.01; ***p < 0.005.

Three independent replicates of SDA analyses with two glass disks for each replicate were performed for the three internal conjugation points. As a result, images of disks were taken with low (2000 rpm) and high (4000 rpm) spinning speeds (**Figure 4C, D**). Bead detachment was only observed at the very outer edge of the disk after spinning at the low speed (**Figure 4C & D**, left). At the higher speed, we found that the central area over which the beads remained attached was larger for N22 and S47 than for S40 (**Figure 4C & D**, right). We used image analysis to calculate and plot the normalized bead density against the shear stress, and fit the sigmoids to a probabilistic model equation to extract final τ50 values for low (**Figure 4E**) and high (**Figure 4F**) spinning speeds. These data demonstrate that when AFF was immobilized through N22 or S47, the beads adhered to the disk with τ50 value of 149 ± 11 and 154 ± 24 dyne cm^-2^ at 2000 rpm and 159 ± 30 and 161 ± 32 dyne cm^-2^ at 4000 rpm, respectively. When AFF was immobilized through S40, we measured lower τ50 values of 129 ± 15 and 125 ± 14 dyne cm^-2^ at 2000 and 4000 rpm, respectively. Normalized τ50 value for each replicate showed statistical significance at both low (**Figure 4G**) and high speeds (**Figure 4H**). The total number of beads remaining on the disk after spinning also showed a consistent trend and similarly reported on the stability of the AFF:(PD-L1) interaction dependent on the anchor point (**Figure 4I, J**). Therefore, the results from the microbead adhesion assays were consistent with AFM-SMFS and SMD simulations for the loading rate range >10^4^ pN s^-1^, where the rupture force of the S47 pulling point was similar to or slightly higher than that of N22, and the rupture force of S40 was lower.

## Discussion

We reported experimental AFM-SMFS analysis, SMD simulations and microbead adhesion assays that demonstrate how altering the surface attachment point (i.e. anchor point) can be used to modulate the mechanical stability of a therapeutic AFF:(PD-L1) complex. Our AFF scaffold is small in size with limited candidate regions eligible for pulling point insertion. We tested 5 pulling positions within AFF and found significant effects on the mechanostability of the AFF:(PD-L1) complex. Depending on the pulling point, these effects included changes in the shape of the unbinding/unfolding energy landscape, different rupture force ranges and loading rate dependencies, along with the occurrence of partially unfolded intermediate states of PD-L1, and catch bond behavior. Computational SMD simulations provided detailed molecular insight into the AFF:(PD-L1) complex with consistent tendencies in rupture force ranges as compared with experimental AFM-SMFS. Based on the parameters extracted from both experimental measurements and simulations, we depicted the energy landscape of the AFF:(PD-L1) complex dissociation reaction as a function of AFF pulling position (**Figure 5A, B**). The steep energy barrier found for N22 contributed to the highest mechanostability by AFM-SMFS analysis. The energy barrier for S47, another mechanostable pulling geometry, was not as high, but the shorter Δx made it the most mechanostable geometry at high loading rates. On the contrary, the longer Δx found for anchor points M1 and S40 made them the least resistant to external force.

**Figure 5.**
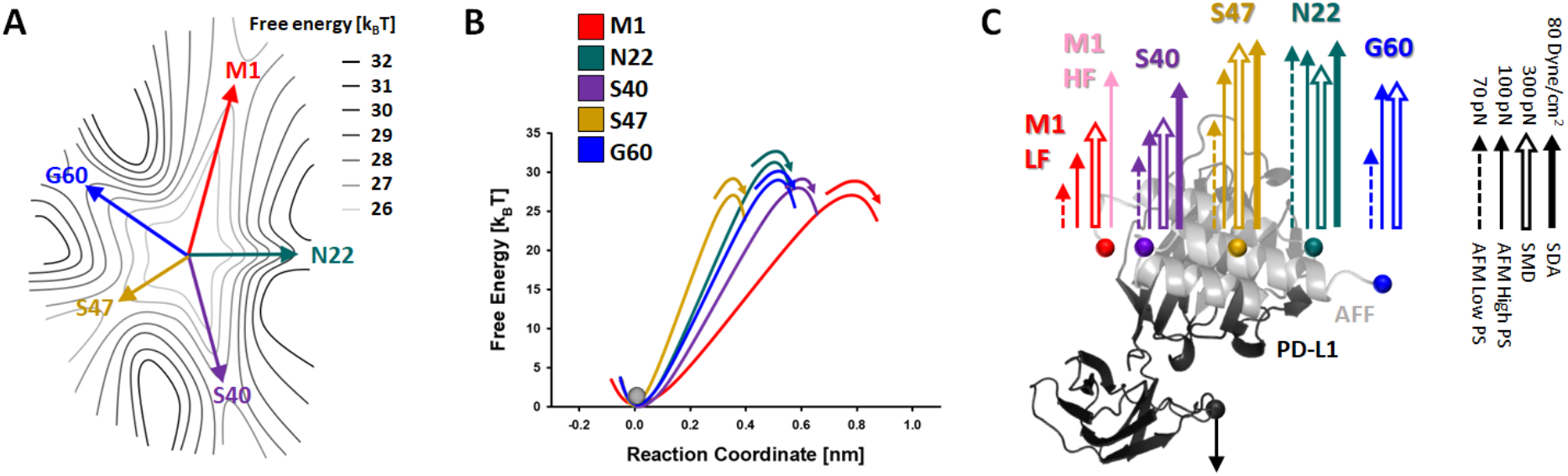
Depictions of mechanical anisotropy of the AFF:(PD-L1) complex. (A) Energy landscape with the five pulling points represented as colored arrows. (B) Unbinding energy barrier heights and barrier positions extracted from DHS model fitting for each pulling geometry. (C) Mechanical anisotropy mapped onto the AFF:(PD-L1) complex structure. The level of mechanostability is indicated by the length of arrows. Dashed arrow: rupture force from AFM-SMFS at the lowest pulling speed, 0.1×10^3^ nm s^−1^; line arrow: rupture force from AFM-SMFS at the highest pulling speed, 6.4×10^3^ nm s^−1^; open arrow: rupture force from SMD at 2.5 × 10^9^ nm s^−1^; filled arrow: τ_50_ from SDA at 4000 rpm. Quantitative scale of arrows is indicated by the black arrows on the right.

We also addressed how these molecular mechanical effects could influence collective and multivalent interactions using microbead adhesion assays under shear stress. This analysis more closely represents a therapeutic or drug delivery scenario for this particular complex, where dynamic interactions and multivalency can influence the response under shear stress. The SDA analyses revealed different τ50 values from three internal pulling points which were consistent with single-molecule analysis by AFM-SMFS and SMD simulations.

We mapped the various measurement and simulation results describing the mechanostability of the AFF:(PD-L1) complex as a function of AFF pulling point onto the protein structure (**Figure 5C**). In all approaches, the analyses were generally consistent and demonstrated significant differences in mechanostability for the various anchor points. Equilibrium binding affinity meanwhile did not change upon introduction of the pulling point mutations, therefore our findings demonstrate how the mechanostability of protein complexes including therapeutic non-antibody scaffolds can be quantified and engineered by altering the loading geometry. This concept suggests that therapeutic efficacy of drug- and nanoparticle targeting proteins can potentially be improved by selecting conjugation points for payloads with optimal stability under force.

## Acknowledgments

This work was supported by the University of Basel, ETH Zurich and a Consolidator Grant from the Swiss State Secretariat for Education, Research and Innovation (SERI) to MN. RCB and DEBG are supported by the National Science Foundation under Grant MCB-2143787, and the National Institute of General Medical Sciences (NIGMS) of NIH through the grant R24-GM145965.

## Author Contributions

B.Y. and M.A.N. conceived the study. B.Y. and Z.L. prepared biological samples. B.Y. performed AFM-SMFS and flow cytometry-based binding experiments, and analyzed the data. B.Y. and M.S.S. performed SDA experiments and analyzed the data. D.E.B.G. performed SMD simulations. R.C.B. and M.A.N. supervised the project. B.Y., D.E.B.G., and M.A.N. drafted and edited the manuscript with input from all authors. All authors reviewed the manuscript.

## Supplementary Information

Electronic supplementary information is available for this article.

