## Supplementary Information for "Engineering the Mechanical Stability of a Therapeutic Affibody/PD-L1 Complex by Anchor Point Selection"

#### Materials & Methods

**Plasmids available on addgene:**
Addgene plasmid #157674: pET28a-ybbR-His-ELP(MV7E2)3-FLN-SpyCatcher

**Cloning of PD-L1-ECD-HIS and PD-L1-ECD-HIS-SpyTag**
DNA sequence of extracellular domain of human PD-L1 was chemically synthesized based on the codon usage of E. coli (GeneArt, Thermo Fisher Scientific) and introduced into pET28a vector via NdeI and XhoI restriction sites generating a new vector pET28a-PD-L1-ECD-HIS, confirmed by further DNA sequencing analysis.

For the AFM-SMFS analysis, SpyTag was further introduced at the C-terminus of PD-L1 by PCR using primer #1 and #2 (Table S1) based on the plasmid pET28a-PD-L1-ECD-HIS and following Gibson assembly with master mix (NEB) generating a new vector pET28a-PD-L1-ECD-HIS-SpyTag, which was confirmed by further DNA sequencing analysis.

**Cloning of AFF-HIS, Fgβ-AFF-HIS, AFF-N-TAG-HIS (M1), AFF-N22TAG-HIS (N22), AFF-S40TAG-HIS (S40), AFF-S47TAG-HIS (S47), and AFF-C-TAG-HIS (G60)**
DNA sequence of Anti-PD-L1 Affibody (AFF) was chemically synthesized based on the codon usage of E. coli (GeneArt, Thermo Fisher Scientific) and introduced into pET28a vector via NdeI and XhoI restriction sites generating a new vector pET28a-Anti-PD-L1-AFF-HIS (for preparation of AFF-HIS), which was confirmed by further DNA sequencing analysis.

For the AFM-SMFS analysis, Fgβ was further introduced at the N-terminus of AFF by PCR using primer #3 and #4 (Table S1) based on the plasmid pET28a-Anti-PD-L1-AFF-HIS and following Gibson assembly with master mix (NEB) generating a new vector pET28a-Fgβ-Anti-PD-L1-AFF-HIS, which was confirmed by further DNA sequencing analysis.

For the incorporation of pAzF into AFF, five different positions were decided (M1, N22, S40, S47, and G60) and amber codon was introduced at each position by site directed mutagenesis using the Q5® Site-Directed Mutagenesis kit (NEB) with primer #5 and #6 for M1, primer #7 and #8 for N22, primer #9 and #10 for S40, primer #11 and #12 for S47, and primer #13 and #14 for G60 (Table S1), generating new plasmids pET28a-Anti-PD-L1-AFF-N-TAG-HIS, pET28a-Anti-PD-L1-AFF-N22TAG-HIS, pET28a-Anti-PD-L1-AFF-S40TAG-HIS, pET28a-Anti-PD-L1-AFF-S47TAG-HIS, and pET28a-Anti-PD-L1-AFF-C-TAG-HIS, which was confirmed by further DNA sequencing analysis.

**Expression, Refolding, and Purification of PD-L1 Variants**
The plasmid with the sequence of PD-L1 variants was introduced into competent E. coli BL21(DE3) strain. Recombinant cells were cultured in 5 ml of Luria-Bertani (LB) medium with 50 μg ml-1 kanamycin at 37 °C overnight. The culture was transferred to 50 mL of Terrific broth (TB) medium with 50 μg ml-1 kanamycin and cultivated at 37 °C and 200 rpm until an optical density at 600 nm (OD600) of ~0.8-1.0 was reached. The expression of recombinant protein was induced by the addition of 1.0 mM isopropyl-β-D-thio-galactopyranoside (IPTG) and the culture was further incubated at 37 °C and 200 rpm for ~9 hrs. The cells were harvested by centrifugation at 4,000 g for 20 min at 4 °C.
The harvested cell pellet was resuspended in a denaturing lysis buffer (10 mM Tris-Cl, and 8M urea; pH ~8). Resuspended cells were placed on ice and disrupted for 15 min using a sonic dismembrator using a 3 s on: 5 s off pattern to allow cooling between each pulse. The lysate was centrifuged at 14,000 g for 20 min at 4 °C. The supernatant was collected and incubated with Ni-NTA resin for 30 min at room temperature to

allow the His6-tagged proteins to bind to the Ni-NTA resin. Then, the mixture was loaded onto a column. The resin was washed with 10–20 resin volumes of wash buffer (20 mM imidazole, 10 mM Tris-Cl, and 8M urea; pH ~8). Recombinant proteins were eluted in the elution buffer (500 mM imidazole, 10 mM Tris-Cl, and 8M urea; pH ~8). The eluted protein solution was serially dialyzed to 8 M, 4 M, 2 M, and 0 M Urea with 5% glycerol, 5% sucrose, 1% arginine, 0.5 mM NaCl in 20 mM Tris-Cl (pH 7.4), and finally to 1x PBS buffer. Precipitation during dialysis was removed by centrifugation at 14,000 g for 20 min at 4 °C and supernatant was further purified by SEC column.

**Expression and Purification of AFF Variants**

The plasmid with the sequence of AFF variants was introduced into a competent E. coli BL21(DE3) strain. Recombinant cells were cultured in 5 ml of Luria-Bertani (LB) medium with 50 μg ml-1 kanamycin at 37 °C overnight. The culture was transferred to 50 mL of Terrific broth (TB) medium with 50 μg ml-1 kanamycin and cultivated at 37 °C and 200 rpm until an optical density at 600 nm (OD600) of ~0.8-1.0 was reached. The expression of recombinant protein was induced by the addition of 1.0 mM isopropyl-β-D-thio-galactopyranoside (IPTG) and the culture was further incubated at 37 °C and 200 rpm for ~9 hrs. The cells were harvested by centrifugation at 4,000 g for 20 min at 4 °C. The cells were harvested by centrifugation at 4,000 g for 20 min at 4 °C. The harvested cell pellet was resuspended in lysis buffer (50 mM Tris, 50 mM NaCl, 0.1% Triton X-100, 5 mM MgCl2; pH 8.0), and disrupted with a sonic dismembrator. The lysate was centrifuged at 14,000 g for 20 min at 4 °C. The supernatant was collected and incubated with Ni-NTA resin, loaded onto a column, washed with wash buffer (1x PBS with 20 mM imidazole; pH 7.4), and eluted in elution buffer (1x PBS with 500 mM imidazole; pH 7.4). The eluted protein solution was further purified by the SEC column.

**Amber Suppression**

Five AFF variants for the free diffusion system of AFM-SMFS analysis were prepared by site-specific incorporation of p-azido-l-phenylalanine (pAzF) at each pulling point using the amber suppression method. The plasmid with the sequence of AFF variants was co-introduced into competent E. coli BL21(DE3) strain with the plasmid pEVOL-pAzF (addgene #31186). The cell culture was transferred to 100 mL of Luria-Bertani (LB) medium with 50 μg ml-1 kanamycin and 34 μg mL-1 chloramphenicol and cultivated at 37 °C and 200 rpm until an optical density at 600 nm (OD600) of ~0.8-1.0 was reached. Then, the cells were collected, washed with ice-cold 0.9% NaCl twice, and transferred to 100 mL M9 medium supplemented with 50 μg mL-1 kanamycin, 34 μg mL-1 chloramphenicol, 1 mM pAzF and 0.02% arabinose. The culture was incubated at 37 °C for 1 h and the expression of recombinant protein was induced by the addition of 1.0 mM IPTG and the culture was further incubated at 20 °C and 200 rpm overnight. Purification of the protein was carried out in the same manner as illustrated previously.

**Conjugation of Fgβ peptide to AFF**

Fgβ-StrepTag-DBCO peptide (JPT Peptide Technologies GmbH, Berlin, Germany) was added to pAzF-incorporated AFF with a molar ratio of 3:1. The mixture was incubated at room temperature with shaking for 1 h, followed by incubation at 4 °C overnight. Successful conjugation was confirmed by SDS-PAGE analysis and conjugated AFF was further purified with SEC in 1x PBS buffer to remove the excess peptide.

**High-resolution mass spectrometry**

Purified AFF variants and Fgβ-conjugated AFF variants protein solution was desalted using Zeba$^{TM}$ spin desalting column (Thermo Fisher Scientific) and diluted to a concentration of 0.2-1.0 mg mL-1 with a final concentration of 0.1% formic acid. The separation of the sample protein analytes was carried out using an UltiMate™ 3000 UHPLC-system equipped with 50 mm Phenomenex Jupiter C4 column (Thermo Fischer Scientific) with a diameter of 2.0 mm, 300 Å pore size, and 5 μm particle size. 1 μL of protein solution was injected for all analyses and the column was kept at 30 °C. HRMS-spectra were acquired on a Bruker maXis

4G ESI-QTOF (Bruker Daltonics) and data deconvolution was done with Bruker Compass DataAnalysis 4.4.

**Surface preparation and protein immobilization for AFM-SMFS**

The surface modification of cantilever and coverglasses and the protein immobilization were done in the same manner as previously illustrated (Figure 1D). Cantilevers were cleaned by UV-ozone treatment for 40 min and cover glasses were soaked in piranha etching solution and rinsed with distilled water (DW). Then, cantilevers and coverglasses were treated with 3-Aminopropyl (diethoxy) methylsilane (APDMES, ABCR GmbH, Karlsruhe, Germany) to silanize the surfaces with amine groups. The amine groups subsequently reacted to a NHS group from sulfosuccinimidyl 4-(N-maleimidomethyl)cyclohexane-1-carboxylate (sulfo-SMCC; Thermo Fischer Scientific) in 50 mM HEPES buffer pH 7.5 for 30 min. The thiol group from Coenzyme A (CoA, 200 µM) reacted to a maleimide group from sulfo-SMCC in coupling buffer (50 mM sodium phosphate, 50 mM NaCl, 10 mM EDTA, pH 7.2) for 2 hrs. Finally, the ybbR-tagged proteins SdrG-FLN-ELP-His-ybbR and pre-conjugated ybbR-His-ELP-FLN-SpyCatcher:PD-L1-SpyTag were site-specifically anchored to the surface using SFP-mediated ligation to CoA in $Mg^{2+}$ supplemented 1x PBS buffer. This resulted in covalent immobilization of SdrG and PD-L1 to cantilever and cover glasses, respectively. Protein-immobilized cantilevers and coverglasses were extensively washed and kept in 1x PBS buffer prior to immediate use. SpyTag-SpyCatcher conjugation of ybbR-His-ELP-FLN-SpyCatcher and PD-L1-ECD-HIS-SpyTag was done by mixing two proteins with the same molar ratio in 1x PBS buffer and pre-incubation for 1 hr prior to ybbR tag ligation.

**AFM-SMFS measurement and data analysis**

Force spectroscopy measurements of PD-L1 and AFF with different pulling geometry were conducted in the same manner as previously illustrated using automated AFM-based SMFS (Force Robot 300, JPK Instruments) with free-diffusion system via SdrG handle. ~ 1 µM of each Fgβ-conjugated AFF variant was added to the measurement buffer between PD-L1 immobilized coverglass and SdrG immobilized cantilever. SMFS data were recorded in 1x PBS buffer at room temperature with constant pulling speeds of $0.1x10^3$, $0.4x10^3$, $1.6x10^3$, $6.4x10^3$ nm $s^{-1}$. Force-extension curves were acquired, filtered and analyzed by a combination of software available on the AFM instrument and custom python scripts. The data traces were filtered by searching for contour length increments that matched the lengths of the fingerprint domains, FLN (≈36 nm). Theoretical contour length increment was calculated based on the equation $\Delta Lc$ = (0.365 nm/AA) × (# AAs in POI) - Lf, where $\Delta Lc$ is expected contour length increment and Lf is end-to-end length of folded protein domain. For FLN, $\Delta Lc$ = 36.9 nm - Lf, where Lf is typically < 5nm. For the Fgβ-AFF (N-terminal pulling geometry), final rupture force was not significantly strong enough to unfold fingerprint domain FLN. Therefore, data traces without FLN unfolding were also selected. For dynamic force spectra, the rupture or unfolding forces vs. loading rate was plotted and median forces and loading rates for each pulling speed were fitted to Bell-Evans model to estimate the effective distance to the transition state (Δx) and the intrinsic dissociation rate or unfolding rate (koff) in the absence of force.

**Conjugation of FAM to AFF**

Fluorescent dye-conjugated AFF solution was prepared by DBCO-azide click chemistry between DBCO-PEG4-FAM5/6 (Jena Bioscience) and pAzF-incorporated AFFs with different 5 positions (AFF-N-pAzF, AFF-N22pAzF, AFF-S40pAzF, AFF-S47pAzF, AFF-C-pAzF). DBCO-PEG4-FAM5/6 was added to pAzF-incorporated AFFs with a molar ratio of 5:1. The mixture was incubated at room temperature with shaking for 1 h, followed by incubation at 4 °C overnight. Unreacted FAM molecules were removed by further purification via His6x-tag and Ni-NTA resin. Successful conjugation was confirmed by SDS-PAGE analysis and conjugated AFF was further purified with SEC in 1x PBS buffer.

**Binding affinity analysis based on flow cytometry**
The binding affinity between AFF variants and PD-L1 was measured using the Attune NxT (Thermo Fisher Scientific) flow cytometer equipped with a 488 nm and a 561 nm laser. First, SpyCatcher was immobilized onto the surface of amine-functionalized PS beads via ybbR Tag. Then, SpyCatcher-immobilized PS bead was incubated with 10 μM PD-L1-SpyTag solution for > ~2 hrs at RT to conjugate PD-L1 to PS bead by isopeptide bond formation between SpyTag and SpyCatcher. PD-L1 immobilized beads were incubated in FAM-labeled AFF variants' solution with different concentrations ranging from 0.04 nM to 625 nM for ~1-2 hrs at RT. After washing, shift of fluorescence from FAM was recorded and plotted against the concentration of AFF to derive the dissociation constant between PD-L1-S and AFFs with different conjugation geometry.

**Computational models**
The initial model was obtained previously described[47]. Briefly, the sequences for the mature domain of PD-L1 (UniProt Q9NZQ7, 18-234) and the Affibody (1-60) were used as input for prediction using AlphaFold version 2.3.2[49,50] in multimer mode using all 5 available v3 multimer parameter sets, resulting in a total of 25 predictions per sequence pair. The QwikFold VMD's[51] plugin was used to set the experiments and post-process the results, and calculations were run using the Cybershuttle[52] Research Environment deployed at the SDSC Expanse[53] supercomputer.

***Molecular dynamics (MD) simulation***
The (PD-L1):AFFibody were subjected to refinement and conformational sampling by molecular dynamics simulations. The system was then solvated with TIP3P[54] water and neutralized using Sodium atoms as counter-ions, which were randomly arranged in the solvent. Total system sizes were approximately 100k-125k atoms. The MD simulations were performed employing the GPU-accelerated molecular dynamics package NAMD3[55]. The CHARMM36 force field [56,57] was used to describe all systems. The simulations were carried out assuming periodic boundary conditions in the NpT ensemble with temperature maintained at 300 K using Langevin dynamics for pressure, kept at 1 bar, and temperature coupling. A distance cut-off of 12.0 Å was applied to short-range, non-bonded interactions, whereas long-range electrostatic interactions were treated using the particle-mesh Ewald (PME)[58] method. The equations of motion were integrated using the r-RESPA multiple time step scheme[59] to update the van der Waals interactions every step and electrostatic interactions every two steps. The time step of integration was chosen to be 4 fs for all simulations performed. Before the production SMD simulations all the systems were submitted to an energy minimization protocol for 5,000 steps followed by MD simulations with position restraints in the protein backbone atoms were performed for 1.0 ns, progressively raising the temperature from 10K to 300K. These steps were followed by an unrestrained equilibration for additional 5 ns.

***Steered molecular dynamics (SMD) simulation***
In all simulations, totaling over 300 SMD simulations, SMD was employed by harmonically restraining the position of a C-terminal amino acid residue of PD-L1, and moving a second restraint point at constant speed ($2.5x10^7$ , $2.5x10^8$, $2.5x10^9$ nm $s^{-1}$) from each pulling point (Affibody residues M1, N22, S40, S47, and G60), with and extension between 6 (N22, S40, S47) and 10 ($C_{terminal}$,$N_{terminal}$) nanometers, in 40-48 production SMD runs.

***Molecular Dynamics Simulations Analysis***
All analyses of MD trajectories were carried out employing VMD, its plugins and TCL scripts, unless stated differently. Analysis outputs were post-processed to generate graphs using Python3 libraries, including Matplotlib[60], Pandas[61], and Seaborn[62], unless stated differently. Figure panels were assembled with CorelDraw Graphics Suite 2021.

**Cys-SpyCatcher preparation**

Cys-SpyCatcher protein was designed to conjugate AFF variants via maleimide-thiol reaction so that AFF variants can have the same ELP linker for spinning disk assay to SMFS analysis. In Cys-SpyCatcher protein construct, native cysteine in FLN was removed and new cysteine was introduced into the short linker region between ddFLN4 and SpyCatcher. This new DNA sequence was amplified via PCR using primer #15 and #16 (Table S1) using the plasmid #157674 (Addgene).

This PCR-amplified DNA string was assembled with PCR-amplified DNA backbone (via PCR using primers #17 and #18 (Table S1) using the plasmid #157674 (Addgene)) into a new vector pET28a-ybbR-HIS-ELP-ddFLN4-Cys-SpyCatcher using Gibson assembly master mix (NEB) and was confirmed by DNA sequencing.

Newly constructed recombinant plasmid pET28a-ybbR-HIS-ELP-ddFLN4-Cys-SpyCatcher was introduced into a competent E. coli BL21(DE3) strain. Recombinant cells were cultured in 5 ml of Luria-Bertani (LB) medium with 50 µg ml-1 kanamycin at 37 °C overnight. The culture was transferred to 50 mL of LB medium with 50 µg ml-1 kanamycin and cultivated at 37 °C and 200 rpm until an optical density at 600 nm (OD600) of ~0.7 was reached. Then, the expression was induced by the addition of 0.5 mM IPTG and the culture was further incubated at 20 °C and 200 rpm for ~18 hrs. The cells were harvested by centrifugation at 4,000 g for 20 min at 4 °C. The harvested cell pellet was resuspended in lysis buffer (50 mM Tris, 50 mM NaCl, 0.1% Triton X-100, 5 mM MgCl2; pH 8.0). Resuspended cells were placed on ice and disrupted for 15 min using a sonic dismembrator using a 3 s on: 5 s off pattern to allow cooling between each pulse. The lysate was centrifuged at 14,000 g for 20 min at 4 °C. The supernatant was collected and incubated with Ni-NTA resin for 30 min at room temperature to allow the His6x-tagged proteins to bind to the Ni-NTA resin. Then, the mixture was loaded onto a column. The resin was washed with 10–20 resin volumes of wash buffer (20 mM imidazole in 1x PBS). Recombinant proteins were eluted in the elution buffer (500 mM imidazole in 1x PBS). The eluted protein solution was directly treated with DTT and further purified by the SEC column.

**Conjugation of Cys-SpyCatcher to AFF**

Cys-SpyCatcher was DBCO-functionalized using DBCO-PEG4-maleimide (BroadPharm) with reaction between free thiol of Cys-SpyCatcher and maleimide group. SEC-purified Cys-SpyCatcher after DTT treatment was mixed to DBCO-PEG4-maleimide with a ratio of ~1:15-20 at 4 °C, overnight. The following day, DBCO-functionalized Cys-SpyCatcher was purified by the SEC column. After that, AFF variants, AFF-N22pAzF, AFF-S40pAzF, and AFF-S47pAzF, were conjugated to DBCO-Cys-SpyCatcher by mixing between them with a ratio of 1:3 at 4 °C, overnight. Finally, further SEC purification resulted in AFF-pAzF-DBCO-Cys-SpyCatcher variants stock solution.

**Spinning disk assay (SDA) analysis**

The surface modification of coverglasses and the protein immobilization were done in the same manner as previously illustrated. The amine groups of aminosilanized cover glasses reacted to a NHS group from sulfo-SMCC (Thermo Fischer Scientific) in 50 mM HEPES buffer pH 7.5 for 30 min. The thiol group from Coenzyme A (CoA, 200 µM) reacted to a maleimide group from sulfo-SMCC in coupling buffer (50 mM sodium phosphate, 50 mM NaCl, 10 mM EDTA, pH 7.2) for 2 hrs. Finally, each ybbR-tagged AFF-pAzF-DBCO-Cys-SpyCatcher protein variants (AFF-N22pAzF, AFF-S40pAzF, and AFF-S47pAzF) were site-specifically anchored to the surface using SFP-mediated ligation to CoA in Mg2+ supplemented 1x PBS buffer for 2 hrs. This resulted in covalent immobilization of AFF-pAzF-DBCO-Cys-SpyCatcher protein variants to cover glasses for SDA. To reduce the non-specific interaction, protein-immobilized coverglasses were treated with 5% BSA solution for 10 min. The surface modification of PS beads and the PD-L1 immobilization were also done in the same manner.

Then, PD-L1 immobilized PS beads were seeded on the surface in 1x PBS containing 0.1% BSA and allowed to adhere for 30 min at room temperature. Before spinning, the bead suspension was removed and gently replaced with PBS. The coverglasses were mounted on the spinning disk device, secured by vacuum suction, and immersed in a solution of PBS at room temperature. The coverglasses were spun with speeds of 2000 rpm and 4000 rpm for 5 min, imaged by frame at 10× magnification on an Olympus IX81 microscope (~500 individual images automatically stitched together with CellSens software (version 1.16; Olympus, Tokyo, Japan)), and saved. Taken images were further analyzed as illustrated previously with a custom Python-based image analysis script and MATLAB 2022b (The MathWorks, Natick, MA) to extract the fraction of adherent beads at different positions on the disk by normalizing the density of beads at each section of the disk with the density of beads at the center of the disk, where the shear forces are close to zero.[35] The fraction of adherent beads (f) was plotted along with the shear stress (τ; Pa) and fitted to a sigmoid probabilistic model:

$$f = \frac{a}{1 + e^{b(\tau - \tau_{50})}}$$

to extract $\tau_{50}$ value which is the shear stress at which 50% of the beads remain adherent.

#### Protein Sequences

**PD-L1-ECD-HIS**

MFTVTVPKDLYVVEYGSNMTIECKFPVEKQLDLAALIVYWEMEDKNIIQFVHGEEDLKVQHSSYRQRARLL
KDQLSLGNAALQITDVKLQDAGVYRCMISYGGADYKRITVKVNAPYNKINQRILVVDPVTSEHELTCQAEG
YPKAEVIWTSSDHQVLSGKTTTTNSKREEKLFNVTSTLRINTTTNEIFYCTFRRLDPEENHTAELVIPELPLA
HPPNERGSHHHHHH

**PD-L1-ECD-HIS-SpyTag**

MFTVTVPKDLYVVEYGSNMTIECKFPVEKQLDLAALIVYWEMEDKNIIQFVHGEEDLKVQHSSYRQRARLL
KDQLSLGNAALQITDVKLQDAGVYRCMISYGGADYKRITVKVNAPYNKINQRILVVDPVTSEHELTCQAEG
YPKAEVIWTSSDHQVLSGKTTTTNSKREEKLFNVTSTLRINTTTNEIFYCTFRRLDPEENHTAELVIPELPLA
HPPNERGSHHHHHHGSAHIVMVDAYKPTK

**AFF-HIS**

MVDAKYAKERNKAAYEILYLPNLTNAQKWAFIWKLDDDPSQSSELLSEAKKLNDSQAPKGSHHHHHH

**Fgβ-AFF-HIS**

MNEEGFFSARGHRPLDGSGSVDAKYAKERNKAAYEILYLPNLTNAQKWAFIWKLDDDPSQSSELLSEAK
KLNDSQAPKGSHHHHHH

**AFF-N-TAG-HIS (M1)**

MGSGSGS-pAzF-
VDAKYAKERNKAAYEILYLPNLTNAQKWAFIWKLDDDPSQSSELLSEAKKLNDSQAPKGSHHHHHH

**AFF-N22TAG-HIS (N22)**

MVDAKYAKERNKAAYEILYLP-pAzF-
LTNAQKWAFIWKLDDDPSQSSELLSEAKKLNDSQAPKGSHHHHHH

**AFF-S40TAG-HIS (S40)**

MVDAKYAKERNKAAYEILYLPNLTNAQKWAFIWKLDDDP-pAzF-
QSSELLSEAKKLNDSQAPKGSHHHHHH

**AFF-S47TAG-HIS (S47)**

MVDAKYAKERNKAAYEILYLPNLTNAQKWAFIWKLDDDPSQSSELL-pAzF-
EAKKLNDSQAPKGSHHHHHH

**AFF-C-TAG-HIS (G60)**

MVDAKYAKERNKAAYEILYLPNLTNAQKWAFIWKLDDDPSQSSELLSEAKKLNDSQAPKGS-pAzF-
GSGSHHHHHH

**SdrG-FLN-ELP-HIS-ybbR**

MGTEQGSNVNHLIKVTDQSITEGYDDSDGIIKAHDAENLIYDVTFEVDDKVKSGDTMTVNIDKNTVPSDLT
DSFAIPKIKDNSGEIIATGTYDNTNKQITYTFTDYVDKYENIKAHLKLTSYIDKSKVPNNNTKLDVEYKTALSS
VNKTITVEYQKPNENRTANLQSMFTNIDTKNHTVEQTIYINPLRYSAKETNVNISGNGDEGSTIIDDSTIIKVY
KVGDNQNLPDSNRIYDYSEYEDVTNDDYAQLGNNNDVNINFGNIDSPYIIKVISKYDPNKDDYTTIQQTVT
MQTTINEYTGEFRTASYDNTIAFSTSSGQGQGDLPPEGSGSGSGSADPEKSYAEGPGLDGGECFQPSK
FKIHAVDPDGVHRTDGGDGFVVTIEGPAPVDPVMVDNGDGTYDVEFEPKEAGDYVINLTLDGDNVNGFP
KTVTVKPAPGSGSGSHGVGVPGMGVPGVGVPGVGVPGVGVPGVGVPGVGVPGVGVPGVGVPGEGVP
GEGVPGVGVPGMGVPGVGVPGVGVPGVGVPGVGVPGVGVPGVGVPGVGVPGEGVPGEGVPGVGVP
GMGVPGVGVPGVGVPGVGVPGVGVPGVGVPGVGVPGVGVPGVGVPGEGVPGEGVPGWRGHHHHHHGSDSL
EFIASKLA

**ybbR-HIS-ELP-FLN-SpyCatcher**

MGTDSLEFIASKLAHHHHHHWGSGHGVGVPGMGVPGVGVPGVGVPGVGVPGVGVPGVGVPGVGVPG
VGVPGEGVPGEGVPGVGVPGMGVPGVGVPGVGVPGVGVPGVGVPGVGVPGVGVPGVGVPGEGVPG
EGVPGVGVPGMGVPGVGVPGVGVPGVGVPGVGVPGVGVPGVGVPGVGVPGEGVPGEGVPGWPSGS
ADPEKSYAEGPGLDGGECFQPSKFKIHAVDPDGVHRTDGGDGFVVTIEGPAPVDPVMVDNGDGTYDVE
FEPKEAGDYVINLTLDGDNVNGFPKTVTVKPAPGSGSGSGSVDTLSGLSSEQGQSGDMTIEEDSATHIKF
SKRDEDGKELAGATMELRDSSGKTISTWISDGQVKDFYLYPGKYTFVETAAPDGYEVATAITFTVNEQGQ
VTVNGKATKGDAHI

**ybbR-HIS-ELP-FLN-Cys-SpyCatcher**

MGTDSLEFIASKLAHHHHHHWGSGHGVGVPGMGVPGVGVPGVGVPGVGVPGVGVPGVGVPGVGVPG
VGVPGEGVPGEGVPGVGVPGMGVPGVGVPGVGVPGVGVPGVGVPGVGVPGVGVPGVGVPGEGVPG
EGVPGVGVPGMGVPGVGVPGVGVPGVGVPGVGVPGVGVPGVGVPGVGVPGEGVPGEGVPGWPSGS
ADPEKSYAEGPGLDGGESFQPSKFKIHAVDPDGVHRTDGGDGFVVTIEGPAPVDPVMVDNGDGTYDVE
FEPKEAGDYVINLTLDGDNVNGFPKTVTVKPAPGSGSGCGSVDTLSGLSSEQGQSGDMTIEEDSATHIKF
SKRDEDGKELAGATMELRDSSGKTISTWISDGQVKDFYLYPGKYTFVETAAPDGYEVATAITFTVNEQGQ
VTVNGKATKGDAHI

### Supporting Figures & Tables

**Table S1.** Primers

| Number | Sequence |
|---|---|
| 1 | 5'-GTCATGGTTGATGCATACAAGCCGACGAAGTAACTCGAGTAAGATCCGGCTG-3' |
| 2 | 5'-GGCTTGTATGCATCAACCATGACAATATGGGCGCTACCGTGATGATGATGGTGATGGCTAC-3' |
| 3 | 5'-GCTGGATGGTAGCGGTAGCGTGGACGCCAAATATGCCAAAG-3' |
| 4 | 5'-GCTACCGCTACCATCCAGCGGACGATGACCACGTGCGCTAAAAAAGCCTTCTTCATTCATATGTATATCTCCTTCTTAAAGTTAAAC-3' |
| 5 | 5'-AGCGGCTCTGGTAGCTAGGTGGACGCCAAATATGCCAAAG-3' |
| 6 | 5'-GCTACCAGAGCCGCTACCCATATGTATATCTCCTTCTTAAAGTTAAAC-3' |
| 7 | 5'-GTATCTGCCGTAGCTGACCAATG-3' |
| 8 | 5'-AGAATTTCGTAGGCTGCTTTATTAC-3' |
| 9 | 5'-TGATGATCCGTAGCAGAGCAGCGAAC-3' |
| 10 | 5'-TCCAGTTTCCAGATAAATG-3' |
| 11 | 5'-CGAACTGCTGTAGGAAGCAAAAAAACTGAATGATAGCC-3' |
| 12 | 5'-CTGCTCTGGCTCGGATCA-3' |
| 13 | 5'-CTAGGGCTCTGGTAGCCATCACCATCATCATCAT-3' |
| 14 | 5'-GCTACCAGAGCCCTAGCTACCTTTCGGTGCCTGG-3' |
| 15 | 5'-AGCTTCCAGCCGTCTAAATTCAAA-3' |
| 16 | 5'-TCAGGGTATCAACGCTACCGCAACCAGAACCGGAGCCCG-3' |
| 17 | 5'-GGTAGCGTTGATACCCTGAGC-3' |
| 18 | 5'-AATTTAGACGGCTGGAAGCTTTCACCACCGTCCAGACCC-3' |

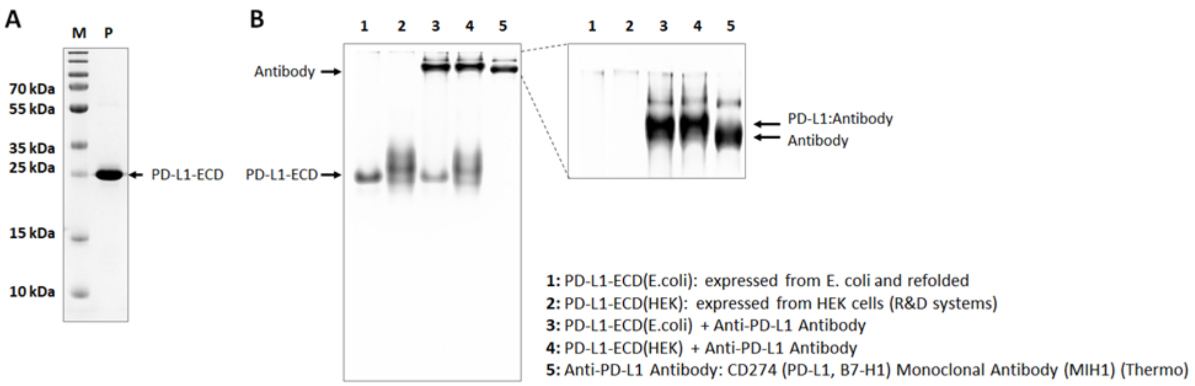


**Figure S1.** Successful expression, refolding, and purification of PD-L1-ECD from E. coli expression system. (A) SDS-PAGE analysis of purified PD-L1-ECD. (B) Native PAGE analysis of purified PD-L1-ECD and its successful binding by anti-PD-L1 antibody which is comparable to binding behavior between commercial recombinant PD-L1-ECD (from HEK cell expression system) and anti-PD-L1 antibody.

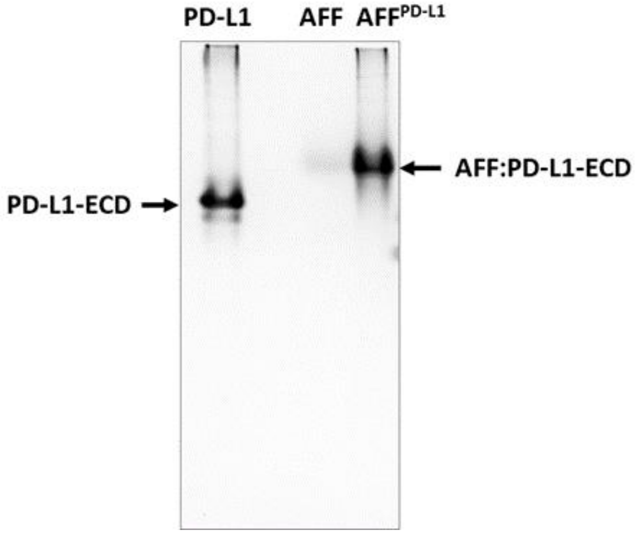


**Figure S2.** Successful preparation of anti-PD-L1 AFF. Binding between purified AFF-HIS and PD-L1-ECD-HIS was confirmed by Native-PAGE analysis.

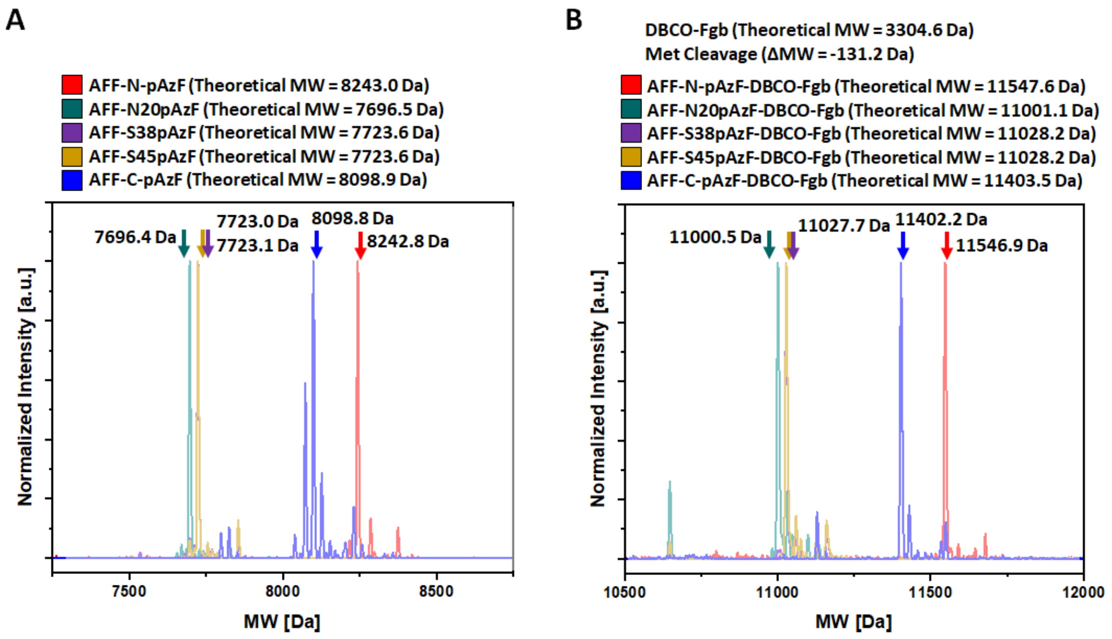


**Figure S3.** HRMS analysis of purified and conjugated AFF-pAzF variants. (A) MW of purified AFF-pAzF variants. (B) MW of purified Fgβ peptide-conjugated AFF-pAzF variants.

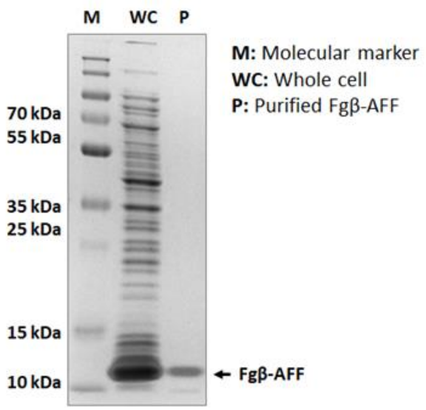


**Figure S4.** Successful preparation of Fgβ-AFF confirmed by SDS-PAGE analysis.

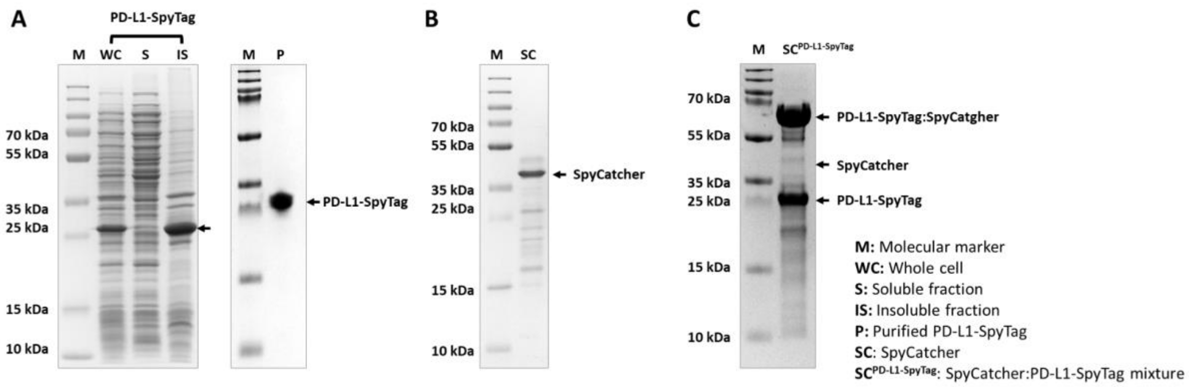


**Figure S5.** Recombinant PD-L1-SpyTag preparation and conjugation to SpyCatcher. (A) Successful expression and purification of PD-L1-SpyTag confirmed by SDS-PAGE analysis. (B) SpyCatcher preparation analyzed by SDS-PAGE. (C) Successful conjugation between PD-L1-SpyTag and SpyCatcher confirmed by SDS-PAGE analysis.

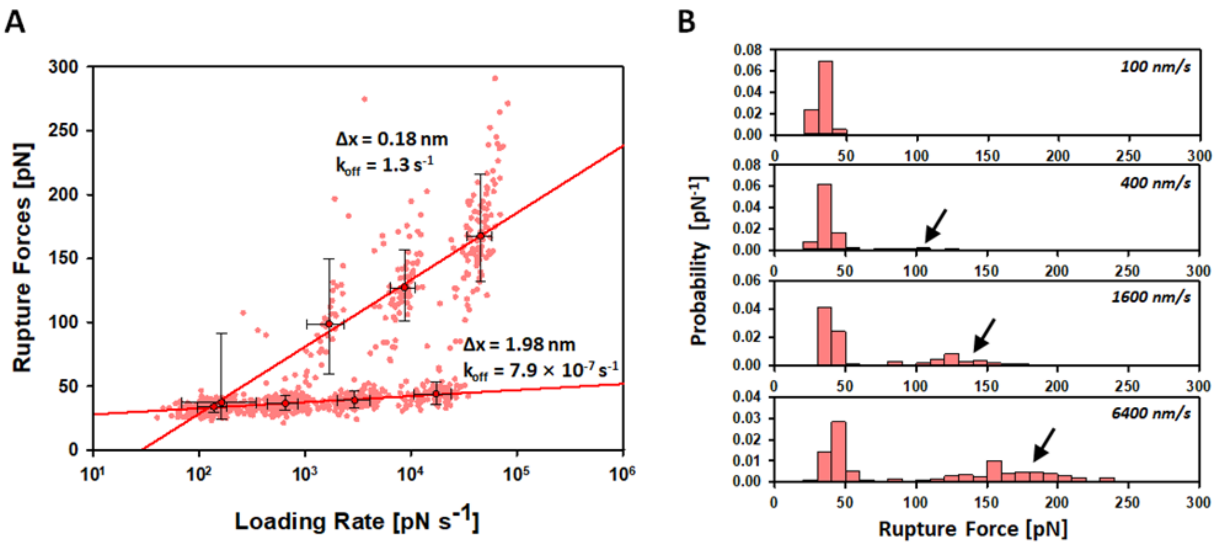


**Figure S6.** Catch bond-like behavior between the Fgβ-AFF:PD-L1 complex rupture. (A) Dynamic force spectra of the Fgβ-AFF:PD-L1 complex rupture. Black-lined circles represent the median rupture force/loading rate at each pulling speed of $0.1 \times 10^3$, $0.4 \times 10^3$, $1.6 \times 10^3$, $6.4 \times 10^3$ nm $s^{-1}$. Error bars are ±1 s.d. Solid lines are least square fits to the Bell-Evans (BE) model. (B) Histograms of the Fgβ-AFF:PD-L1 complex rupture probability at different pulling speeds and emergence of high force population (black arrows).

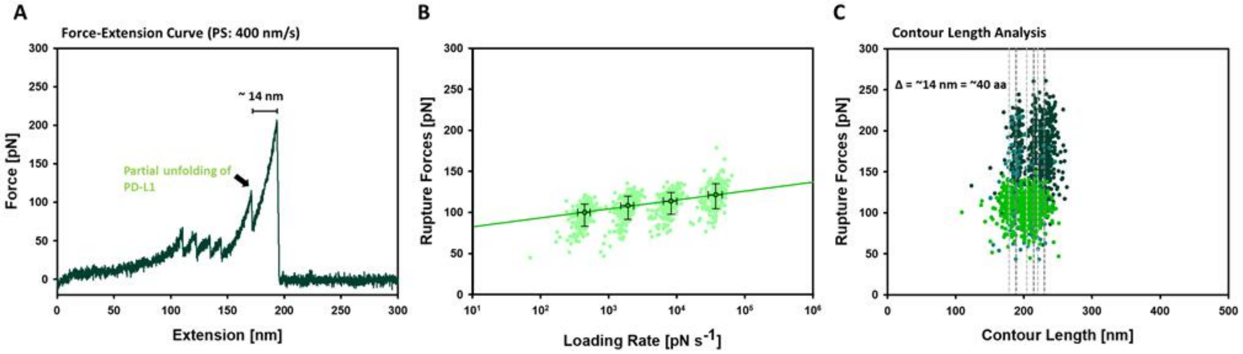


**Figure S7.** Partial unfolding of PD-L1 between the AFF(N22):PD-L1 complex rupture. (A) Typical AFM force−extension traces of the AFF(N22):PD-L1 complex rupture. Additional peak (Lime; partial unfolding of PD-L1) was observed prior to the final rupture between AFF and PD-L1. (B) Dynamic force spectra of the partial PD-L1 unfolding with a pulling geometry of N22. Black-lined circles represent the median unfolding force/loading rate at each pulling speed of $0.1x10^3$, $0.4x10^3$, $1.6x10^3$, $6.4x10^3$ nm $s^{-1}$. Error bars are ±1 s.d. Solid lines are least square fits to the Bell-Evans (BE) model. (C) Contour length analysis showing partial unfolding of PD-L1 as long as ~14 nm which is equivalent to ~40 amino acids long.

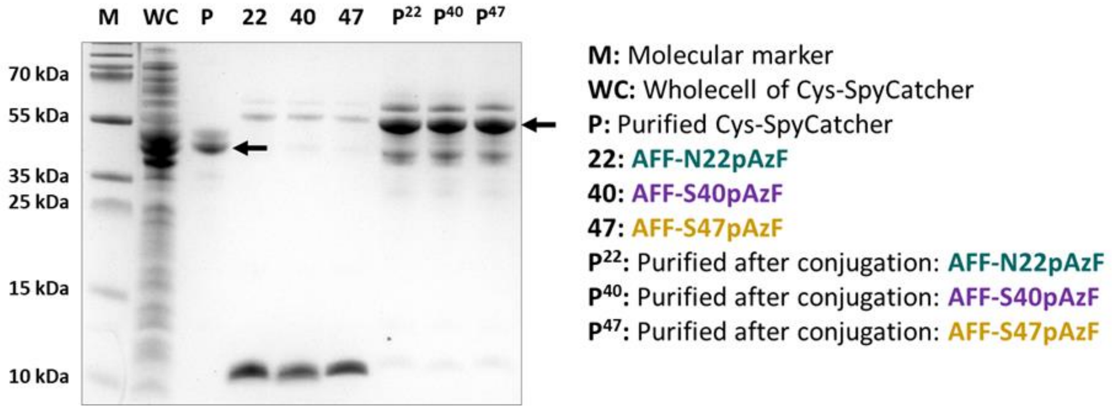


**Figure S8.** Conjugation between Cys-SpyCatcher and AFF-pAzF variants. Successful expression and purification of Cys-SpyCatcher and its following conjugation with AFF-pAzF variants were confirmed by SDS-PAGE analysis.

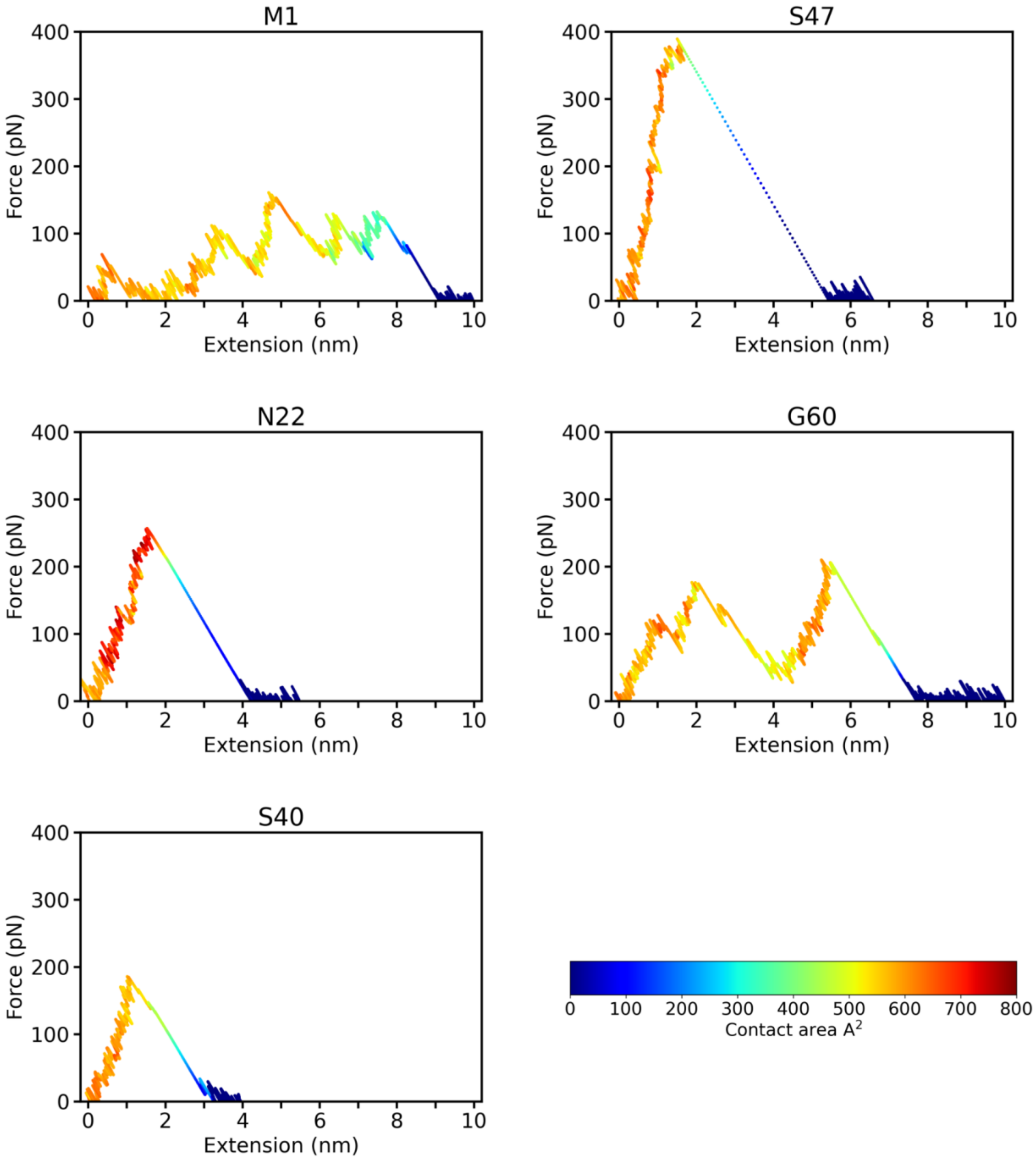


**Figure S9.** Typical force and contact area profiles steered molecular dynamics simulations, pulling at 2.5x10$^7$ nm s$^{-1}$. The dissociation frequently coincides with a sharp loss of contacts area when pulling from the internal residues (N22, S40, and S47), while pulling from the terminal residues (M1 and G60) reveal partial unfolding prior to the final rupture.

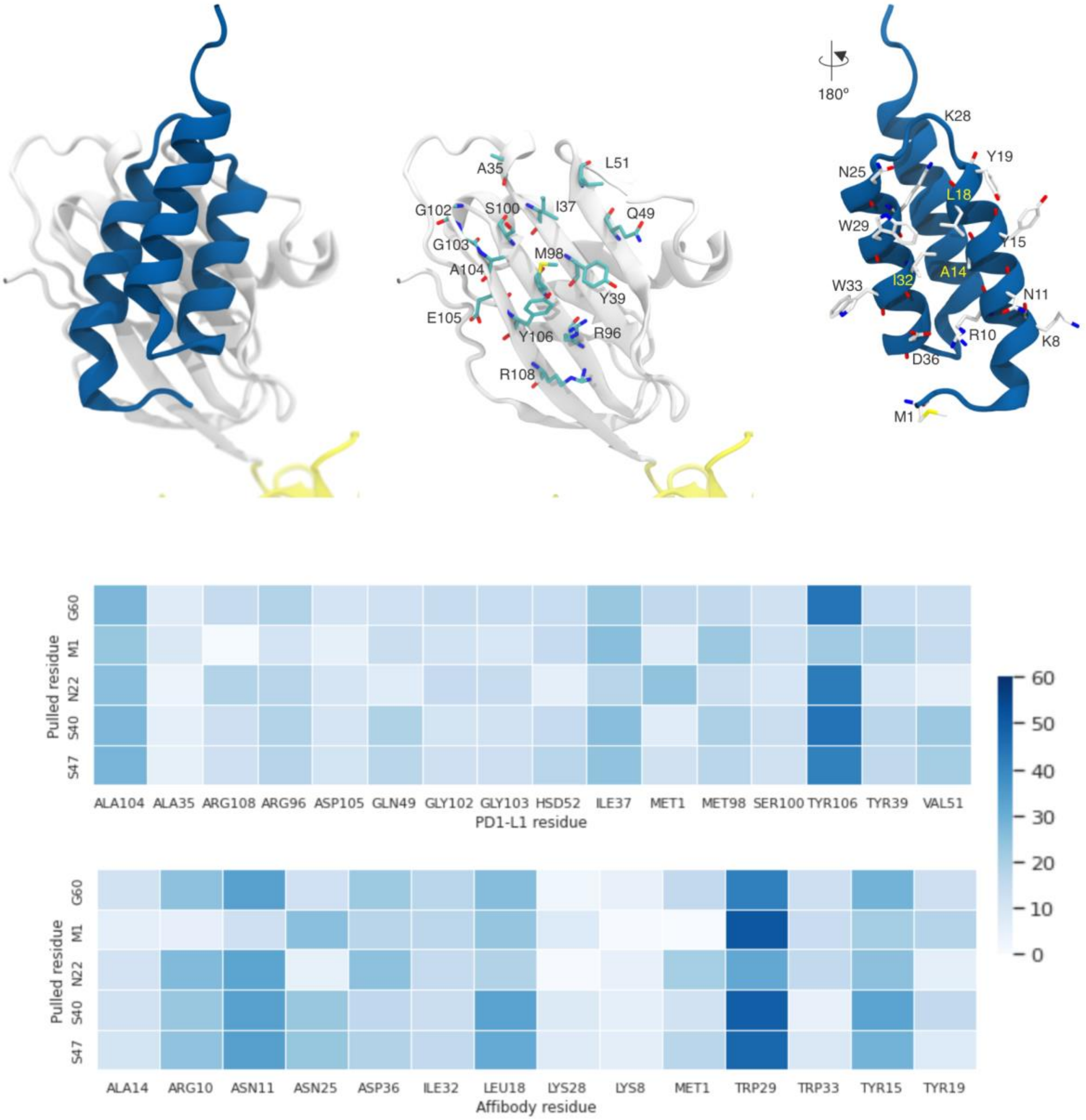


**Figure S10 -** Mean PD1-L1:Affibody contact area (Å$^2$) over 40 replicas for the 5.0 ns before the Peak Rupture Force ($2.5x10^7$ nm $s^{-1}$)

### SI videos

Link to: Video S1
